# GenomeScope 2.0 and Smudgeplots: Reference-free profiling of polyploid genomes

**DOI:** 10.1101/747568

**Authors:** T. Rhyker Ranallo-Benavidez, Kamil S. Jaron, Michael C. Schatz

## Abstract

An important assessment prior to genome assembly and related analyses is genome profiling, where the k-mer frequencies within raw sequencing reads are analyzed to estimate major genome characteristics such as genome size, heterozygosity, and repetitiveness. Here we introduce GenomeScope 2.0 (https://github.com/tbenavi1/genomescope2.0), which applies combinatorial theory to establish a detailed mathematical model of how k-mer frequencies are distributed in heterozygous and polyploid genomes. We describe and evaluate a practical implementation of the polyploid-aware mixture model that, within seconds, accurately infers genome properties across thousands of simulated and eleven real datasets spanning a broad range of complexity. We also present a new method called Smudgeplots (https://github.com/KamilSJaron/smudgeplot) to visualize and infer the ploidy and genome structure of a genome by analyzing heterozygous k-mer pairs. We successfully apply the approach to systems of known variable ploidy levels in the *Meloidogyne* genus and also the extreme case of octoploid *Fragaria x ananassa*.

## 1 Introduction

Genome sequencing has become an integral part of modern molecular biology. The majority of the available analysis methods, however, are designed for established model organisms with chromosome-level reference genomes and detailed annotation readily available. In contrast, genome assemblies of non-model organisms are often fragmented, incomplete, or non-existent. Furthermore, model organisms usually have relatively modest complexity, and are typically haploid or diploid species with relatively low genetic diversity and low repetitive content. Conversely, non-model species often have higher ploidy or higher rates of heterozygosity, and thus are substantially more difficult to analyze. As a result, polyploid species or species with other unusual genome structures are greatly underrepresented among genomics studies.

This underrepresentation reduces the generality of biological insights that can be gleaned from such studies. Notably, polyploids are known to be common, especially among plants and fungi. More than 70% of flowering plants are polyploid (Meyers and Levin 2006) including many crops essential for human consumption and use, including apples, bananas, potatoes, strawberries, and wheat (Renny-Byfield and Wendel 2014). Higher ploidy levels have also been documented in many fungal species (Todd, Forche, and Selmecki 2017). Polyploidy in animals is less common than in these other taxa, but is far from rare, including many species of frogs (Novikova et al. 2019), fish (Comber and Smith 2004), crustaceans and molluscs (Goldman, LoVerde, and Chrisman 1983), as well as many species of nematodes (Szitenberg et al. 2017). The nematode species that are major pests of polyploid crops also happen to be polyploid (Abad et al. 2008). More generally, polyploidization events have important consequences to genome evolution (Otto 2007; Baduel et al. 2018). Developing tools to analyze fragmented and polyploid genomes is therefore essential for our understanding of how polyploidy affects genome and species evolution (Blischak, Kubatko, and Wolfe 2018).

The methods to analyze polyploid genomes typically rely on mapping reads to a haploid reference. However obtaining a complete haploid reference is usually a challenging task (Claros et al. 2012) as the assembly often results in mixed ploidy levels among the assembled sequences depending on the parameter settings (see (Nowell et al. 2018) for an example). Genome assembly has an extra layer of complexity when the basic genomic features of the species are unknown (e.g. size, heterozygosity, and even ploidy). In the context of diploid organisms, several computational approaches have been developed to estimate genome characteristics directly from unassembled sequencing reads, including genome size and heterozygosity (Chikhi and Medvedev 2014; Melsted and Halldórsson 2014; Sun et al. 2018) or repetitiveness and heterozygosity (Simpson 2014). However, none of these approaches model polyploid genomes.

We previously introduced GenomeScope (Vurture et al. 2017), for reference-free analysis of diploid genomes using a statistical analysis of k-mers in unassembled reads, also called the k-mer spectrum. Here we present GenomeScope 2.0, which extends this approach with a polyploid-aware mixture model to computationally infer genome characteristics from unassembled sequencing data. GenomeScope 2.0 fits a mixture of negative binomial distributions to the k-mer spectrum of the sequencing data, with additional components to capture k-mers across higher ploidy levels. To further assist in the analysis of novel species we have also developed Smudgeplot, a visualization technique of genome structure to estimate the ploidy, which is often unknown in non-model organisms. We show that these tools quickly and accurately analyze simulated and real data, including sequencing data from several real genomes (Table S1). These tools can be used to improve the assessment and interpretation of genome assemblies and will substantially aid future studies of polyploid or otherwise complex genomes.

**Table 1:** Mean absolute errors of parameters of 75 simulated triploid datasets, 750 simulated tetraploid datasets, 1,215 simulated pentaploid datasets, and 11,664 simulated hexaploid datasets. Nucleotide divergence refers the the proportion of loci along the polyploid genome for which the nucleotides across all the homologues are not all the same.

| Mean Absolute Errors | Triploid | Tetraploid | Pentaploid | Hexaploid |
| --- | --- | --- | --- | --- |
| Repetitiveness ($d$) | $2.29 \times 10^{-3}$ | $6.61 \times 10^{-3}$ | $9.64 \times 10^{-3}$ | $1.67 \times 10^{-2}$ |
| Nucleotide Divergence | $3.58 \times 10^{-4}$ | $7.38 \times 10^{-4}$ | $1.13 \times 10^{-3}$ | $3.76 \times 10^{-3}$ |
| Monoploid Length | 2,182 bp | 4,320 bp | 5,138 bp | 7,969 bp |

## 2 Methods

### 2.1 Overview

Similar to GenomeScope 1.0 (Vurture et al. 2017), GenomeScope 2.0 takes as input the k-mer spectrum, performs a non-linear least-squares optimization to fit a mixture of negative binomial distributions, and outputs estimates for genome size, repetitiveness, and heterozygosity rates. For example, Figures 1a and 1b show the k-mer profiles, fitted models, and estimated parameters for diploid *Arabidopsis thaliana* and triploid nematode *Meloidogyne enterolobii*. The diploid has two major peaks at approximately 22 and 44, and the triploid has three major peaks centered at approximately 150, 300, and 450. The relative heights of the peaks are proportional to the heterozygosity of the species, and higher coverage peaks represent increasingly higher copy repetitive sequences in the genomes.

**Figure 1:**
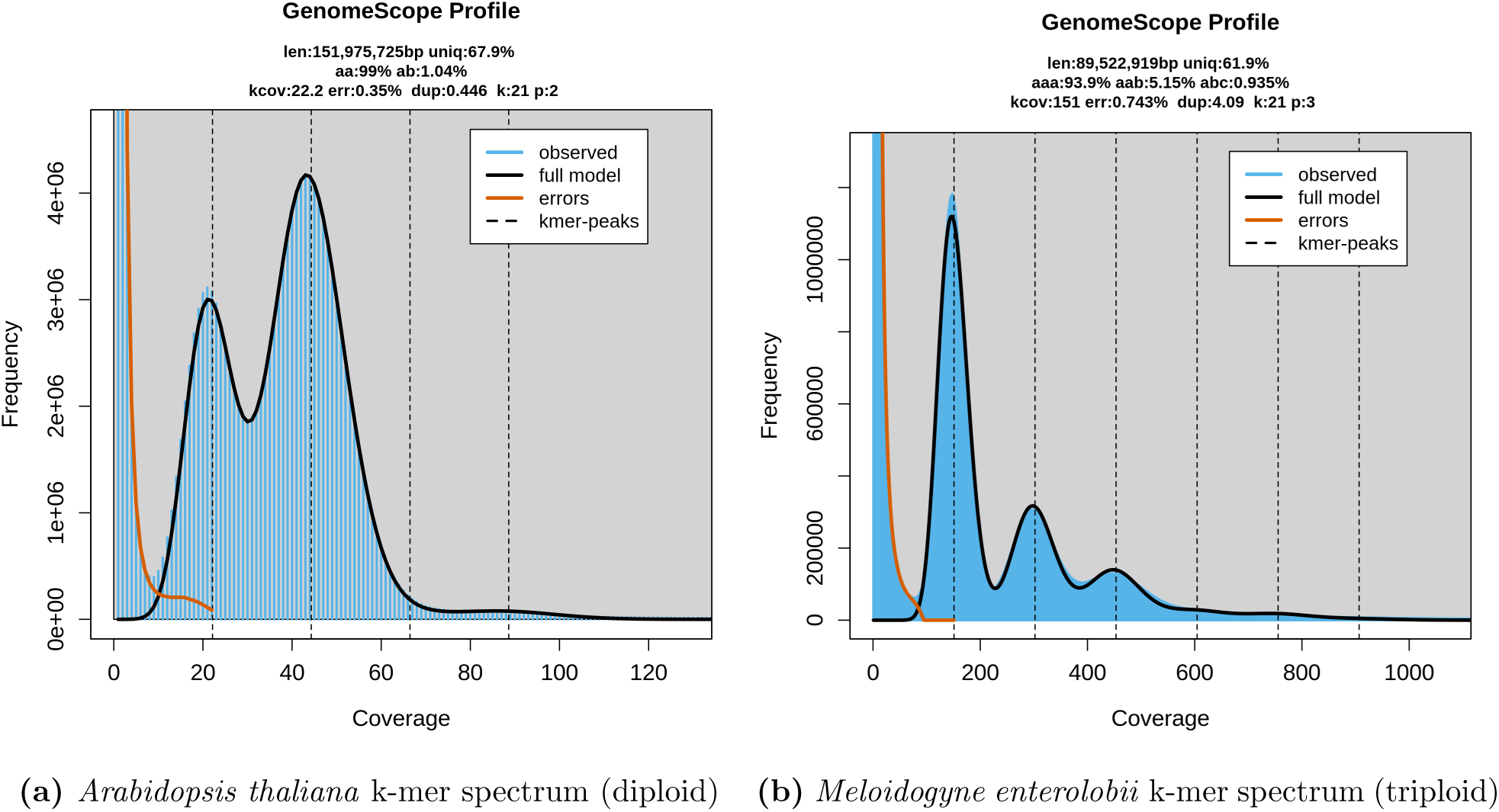
K-mer spectra for representative heterozygous diploid and triploid species. Notice that the diploid plot has two major peaks, while the triploid plot has three. Both also have high frequency putative error k-mers with coverage near 1.

### 2.2 Combinatorial Model

#### 2.2.1 Diploid Model

GenomeScope 1.0 statistically analyzes the k-mer profile and fits a mixture of four negative binomials, the first two representing unique heterozygous and homozygous k-mers, and the next two representing two-copy heterozygous and homozygous k-mers. For example, Figure 1a shows the k-mer profile, fitted model, and estimated parameters for a highly heterozygous diploid *Arabidopsis thaliana* representing an F1 cross between two divergent accessions (Col-0 x Cvi-0, data from (Chin et al. 2016)).

The four negative binomials are equally spaced apart and occur at *λ*, 2*λ*, 3*λ*, and 4*λ* where *λ* = 22.2 is the average k-mer coverage for the diploid genome. More generally, the *i*-th peak corresponds to the contributions from k-mers that occur approximately *i* times in the polyploid genome. It should be noted that although GenomeScope doesn’t fit negative binomials for repetitive regions that occur more than twice, this does not greatly affect the fit on the peaks corresponding to less repetitive regions. This is because the proportion of the genome modeled by a given copy number repeat typically follows a Zeta distribution and hence quickly falls off (Kelley, Schatz, and Salzberg 2010).

The underlying GenomeScope 1.0 model is given by:

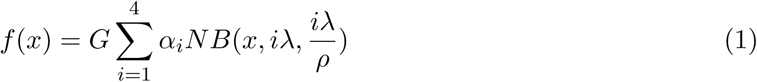

where *f* (*x*) is the k-mer spectrum (i.e. the frequency of the k-mers at coverage depth *x*), *G* is the number of distinct k-mers (i.e. repetitive k-mers are counted only once) in the *monoploid* genome.

Within polyploids, the basic chromosome set from which the other sets are derived, is called the monoploid chromosome set, while the chromosomes present in the gametes of a species constitute the haploid chromosome set. Thus, the monoploid genome consists of a single chromosome set, while the haploid genome typically consists of half of the total number of chromosome sets (Hartl and Jones 1999). Under this model, *α_i_* is, for a single distinct k-mer of the monoploid genome, the expected frequency contribution of the corresponding k-mers across the two homologues to peak *i* of the k-mer spectrum, *NB*(*x, µ, size*) is the negative binomial distribution that approximates the sequencing coverage with mean *µ* and dispersion parameter *size*, *λ* is the average k-mer coverage for the diploid genome, and *ρ* is a bias parameter proportional to PCR duplication and other sequencing biases.

The next crucial step for the model is to mathematically determine the *α_i_* values in terms of the repetitiveness, heterozygosity, and k-mer length. In the diploid case, we have:

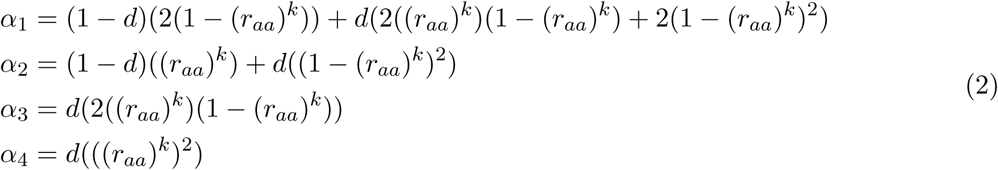

where *d* is the proportion of distinct k-mers of the monoploid genome that occur twice, *r_aa_* is the homozygosity rate, and *k* is the k-mer length.

#### 2.2.2 Polyploid Model

To account for the higher ploidy levels in polyploid organisms, the underlying GenomeScope 2.0 model now fits 2 *× p* negative binomial distributions, where *p* is the ploidy, according to:

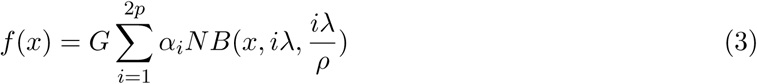

Similarly to the diploid case, each of the 2*p* negative binomials are equally spaced apart and occur at *λ*, 2*λ*, …, and 2*pλ* where *λ* is the average k-mer coverage of the polyploid genome. Again, the *i*-th peak corresponds to the contributions from k-mers that occur approximately *i* times in the polyploid genome.

The next step for the model is to mathematically determine the *α_i_* values in terms of the ploidy, repetitiveness, heterozygosity, and k-mer length. In the polyploid case, this calculation is much more involved and requires utilizing the Möbius inversion formula on partially ordered sets, a classical combinatorics theorem (Rota 1987). For the derivation of this calculation, please refer to Section S1 in the **Online Methods**.

## 3 GenomeScope Implementation

### 3.1 Model fitting algorithm

In order to determine the parameters that best fit the input data, GenomeScope uses a nonlinear least squares minimization technique. While GenomeScope 1.0 used the `nls` function in R based on the Gauss-Newton algorithm, GenomeScope 2.0 instead uses the `nlsLM` function. `nlsLM` utilizes the Levenberg-Marquardt algorithm, with support for lower and upper parameter bounds. Like the Gauss-Newton method, the Levenberg-Marquardt algorithm starts from an initial naive estimate and performs an iterative procedure to update the parameters. However, Levenberg-Marquardt introduces a damping parameter that is adjusted as the iterative process continues, making it more robust. Notable, in many simulated and real datasets with higher ploidy, the `nlsLM` function is able to converge while the `nls` function is not.

### 3.2 Transformed K-mer Histogram

For data sets with high heterozygosity and/or high ploidy the k-mer spectrum does not show clearly defined higher-order peaks. In these cases, fitting to the transformed k-mer spectrum improves the model’s ability to converge. We define the transformed k-mer spectrum as a plot of frequency times coverage (y-axis) versus coverage (x-axis) instead of the typical frequency versus coverage. Transforming the k-mer spectrum effectively increases the heights of higher-order peaks, overcoming the effect of high heterozygosity. This increases the fraction of k-mers in the higher order peaks, especially the homozygous peak, which allows the model to converge. Even for datasets with low heterozygosity and low ploidy, we find fitting to the transformed k-mer spectrum yields accurate results. Consequently, GenomeScope 2.0 now by default fits to the transformed k-mer spectrum. After the fitting process, GenomeScope 2.0 outputs the estimated parameters along with four plots of the best fit model overlaying the k-mer spectrum: 1) untransformed linear, 2) untransformed log, 3) transformed linear, 4) transformed log.

## 4 Smudgeplot

GenomeScope 2.0 is able to accurately analyze organisms given a known ploidy. However, in many cases researchers studying a novel organism may not know the ploidy *a priori*. For this reason, we have implemented Smudgeplots, a new approach to visualize genome structure and infer ploidy directly from the k-mers present in sequencing reads.

For this method, we take as input the set of sequenced k-mers, such as the k-mer frequency files produced by `KMC` (Kokot, Dlugosz, and Deorowicz 2017) or `jellyfish` (Marçais and Kingsford 2011). Then, we search for all pairs of k-mers that differ at exactly one nucleotide through a systematic scan of all input k-mers. To avoid pairing k-mers produced by sequencing errors with genomic k-mers, we search only those k-mers which exceed a coverage threshold (*L*) and assume that such k-mers represent real genomic k-mers. Given how many possible k-mers exist for sufficiently large *k* (e.g. over 4 trillion for *k* = 21), it is very unlikely that two independent genomic k-mers will have the same sequence in all but one nucleotide simply by chance. Thus, the two k-mers in a k-mer pair are homologous and can either represent different alleles of the same locus (heterozygous k-mers) or different loci (paralogs, e.g. duplicated genes or transposable elements). In a reasonably heterozygous genome, the signal from heterozygous k-mers will dominate and therefore can be used to generate an estimate of ploidy.

We denote the two k-mers in each k-mer pair as *A* and *B* such that the coverage of *A* (*CovA*) is always less than or equal to the coverage of *B* (*CovB*). Within every pair, both *A* and *B* can be present in one or more genomic copies and therefore *CovA* + *CovB ∈ {*2*λ,* 3*λ,* 4*λ,* 5*λ, …*, where *λ* is the monoploid genome coverage. Furthermore, the relative minor k-mer coverage 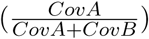 is bounded according to 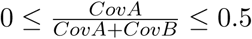 Plotting *CovA* + *CovB* versus 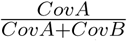 will result in each distinct genomic structure projecting on a different position (i.e. “smudge”) in 2D space (see Figure 2).

**Figure 2:**
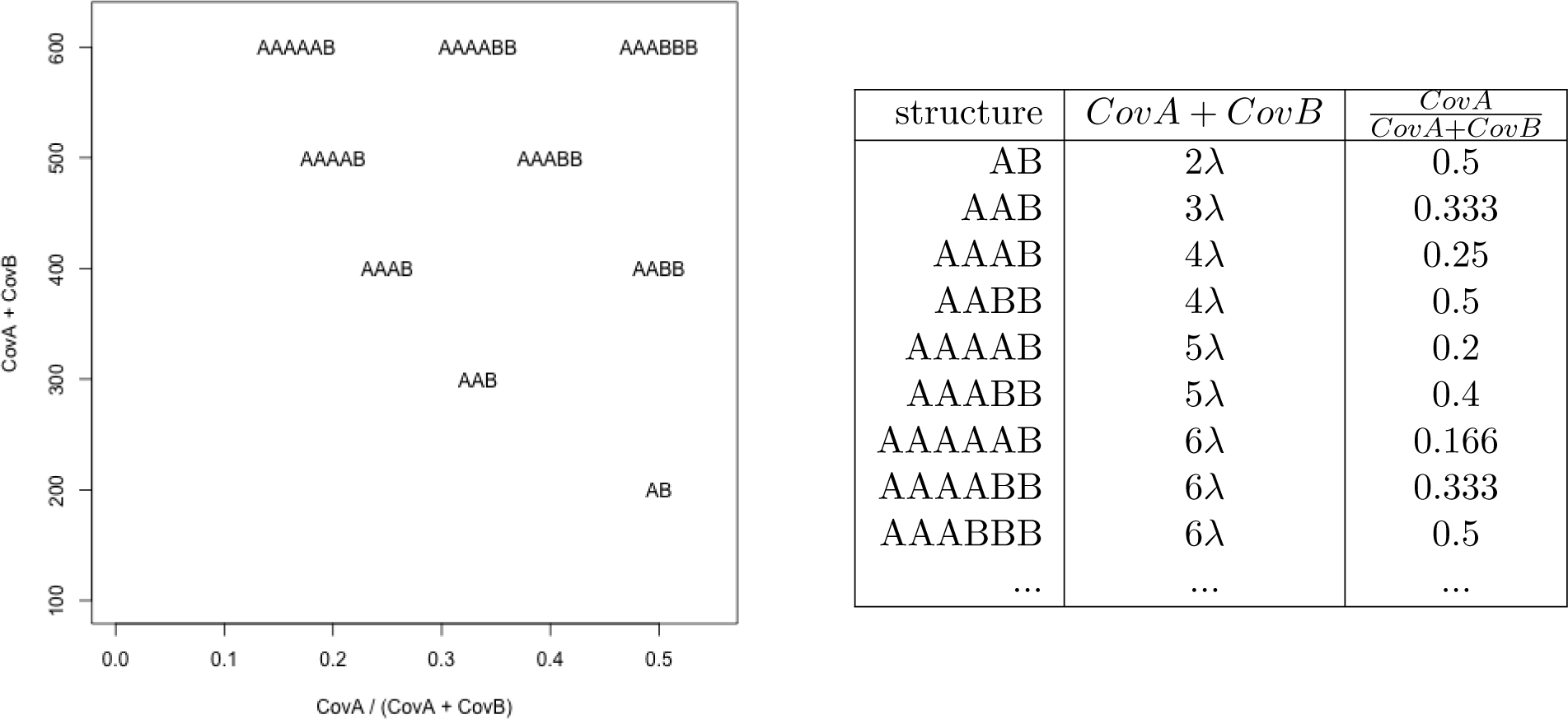
Coordinates of individual genomic structures (for a genome with monoploid coverage (*λ*) equal to 100) in (a) 2D space of coverage sums versus coverage ratios and in (b) a table of coordinates.

By plotting the total coverage of the k-mer pair, *CovA* + *CovB*, versus the relative minor k-mer coverage, 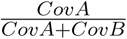, we can identify individual “smudges” that correspond to different haplotype structures. Due to the Poisson nature of the coverages of each position along the genome that is typical in sequencing experiments, the k-mer pairs will not have the exact coordinates as given in Figure 2. However, it is usually possible to resolve the smudge to which each pair belongs. Figure 3a shows an ideal case, where the sequencing coverage is sufficient to completely separate all the smudges, providing very strong evidence of triploidy. The brightness of each smudge is determined by the number of k-mer pairs that fall within it.

**Figure 3:**
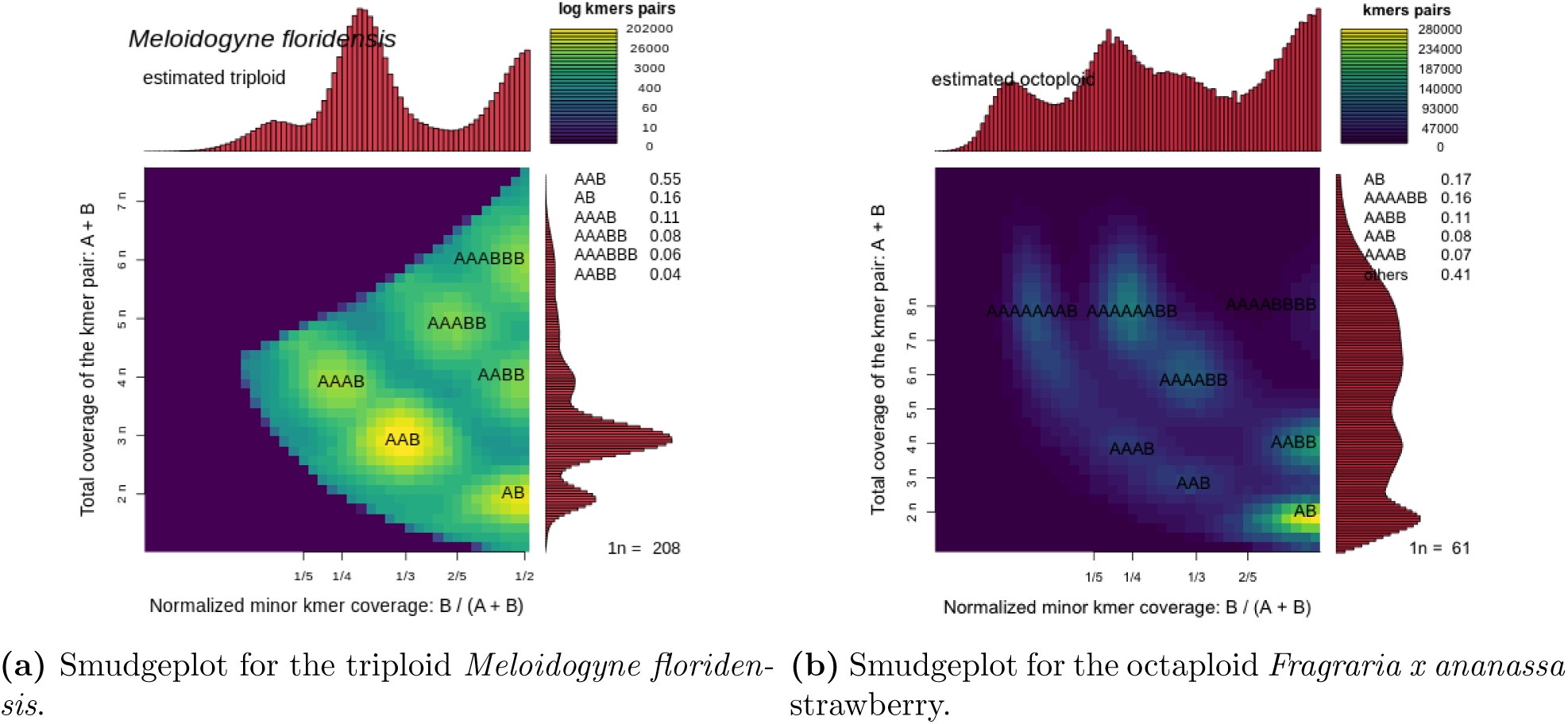
Smudgeplots on real datasets.

The annotation of smudges consist of three steps: 1) identification of smudge boundaries, 2) smudge filtering and 3) estimation of monoploid coverage. First, the 2D space is divided into bins and the number of k-mer pairs in each bin is calculated. Then, the centers of each smudge are chosen to be the bins corresponding to local maxima (in terms of the number of k-mer pairs). The k-mer pairs in all the other bins are aggregated to the nearest neighbouring bin that is designated as a smudge center. Once the boundaries of individual smudges are estimated, we filter smudges that represent less than 0.5% of the data set (i.e. they contain less than 0.5% of the k-mer pairs), as these usually represent repetitive structures of the genome and are frequently misplaced due to too few k-mers representing them.

For the first estimation of the monoploid coverage, we calculate an estimate for each of the identified smudges, and then calculate an overall estimate as the weighted mean of these estimates where the weights are the number of k-mer pairs within each smudge. To calculate the estimate for an individual smudge, we first label the smudge according to its putative structure. For example, of all the smudges with a relative minor coverage near 0.5, the one with the lowest sum of coverages is assumed to be AB and others are labeled using the AB smudge as a reference. This process is continued for all relative minor coverages of the identified smudges until all smudges are labeled. Finally, the estimate of monoploid coverage for an individual smudge is given by its sum of coverages divided by the number of k-mers that make up its labeled structure. For example, the estimate for an AAB smudge would be 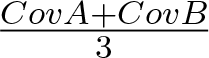 since three k-mers make up the AAB structure.

Next, this first estimate of monoploid coverage is used to re-annotate smudges and subsequently to estimate the ploidy. If multiple smudges get annotated with the same genome structure, the whole process is repeated with lowered resolution (i.e. the number of bins in the 2D plot is decreased). This estimate of monoploid coverage assumes that we correctly labeled each smudge with its putative structure, which may not be the case if we didn’t correctly find the smudge with lowest sum of coverages for a given relative minor coverage. Therefore, the final estimate of monoploid coverage is refined by using kernel smoothing applied on the subset of k-mer pairs within the brightest smudge in the Smudgeplot.

The Smudgeplot estimates of monoploid coverage (*λ*) and ploidy allow users to visualize and discover properties about genomes with high levels of imperfect duplications, various ploidy levels, and high heterozygosity. This visualization tool is especially powerful in combination with GenomeScope, as both independently estimate monoploid coverage by exploiting different genomic properties. Notably, Smudgeplot is able to accurately predict that *Fragraria x ananassa* is octaploid (see Figure 3b).

## 5 Results

### 5.1 Simulated Polyploid Genomes

We first applied GenomeScope 2.0 on 13,704 simulated datasets with varying ploidy (3, 4, 5, and 6), repetitiveness (0%, 10%, and 20%), and nucleotide heterozygosity rates (0%, 0.5%, 1%, %, and 2% for ploidies 3 and 4; 0%, 1%, and 2% for ploidies 5 and 6). For each ploidy, we also simulated all the possible topological relationships between the homologous chromosomes. For example, for tetraploid organisms there are two possible topologies: *AAAA → AAAB → AABC → ABCD* which corresponds to an autotetraploid topology and *AAAA → AABB → AABC → ABCD* which corresponds to an allotetraploid topology (see Section S2 in the **Online Methods** for further explanation). For pentaploid organisms there are five possible topologies, and for hexaploid organisms there are sixteen possible topologies.

Each triploid topology consists of two nucleotide heterozygosity forms (e.g. *aab* and *abc*), while each tetraploid, pentaploid, and hexaploid topology consists of three, four, and five heterozygosity forms respectively. Thus, we simulated 75 triploid datasets (3 repetitiveness values, 5 heterozygosity values for each of the 2 heterozygosity forms, 1 topology), 750 tetraploid datasets (3 repetitiveness values, 5 heterozygosity values for each of the 3 heterozygosity forms, 2 topologies), 1,215 pentaploid datasets (3 repetiveness values, 3 heterozygosity values for each of the 4 heterozygosity forms, 5 topologies), and 11,664 hexaploid datasets (3 repetiveness values, 3 heterozygosity values for each of the 5 heterozygosity forms, 16 topologies).

For the simulated data, we simulated 15x coverage per homologue and 1% sequencing error, to test GenomeScope 2.0 in relatively poor data quality conditions. Each simulated dataset was created with a generative model using a random 1 Mbp monoploid genome as a “progenitor.” To test GenomeScope’s robustness on genomes of varying size, we also simulated using progenitor genomes of size 1 Mbp, 10 Mbp, 100 Mbp, and 1 Gbp. The mean absolute errors of the estimated parameters on the simulated datasets are shown below, which demonstrate that GenomeScope 2.0 is highly accurate. For the full results, see the **Supplemental Files**.

We then performed more specific testing to validate GenomeScope 2.0’s performance at predicting nucleotide divergence, repetitiveness, and length. Specifically, for each of these three parameters, we held the others constant, and varied only the parameter of interest:

- For nucleotide divergence, we systematically evaluated across 0% to 25% in 0.5% increments, for a total of 51 values. We used a 100 Mbp progenitor genome, 15x coverage per homologue, and 1.0% sequencing error. Figure 4 below shows the difference between the estimated and true nucleotide divergence as a function of the true nucleotide divergence, for ploidies 3, 4, 5, and 6.
- For repetitiveness, we evaluated a parameter sweep from 0% to 50% in 1% increments, for a total of 51 values. We used a 100 Mbp progenitor genome, 15x coverage per homologue, and 1.0% sequencing error. Figure 5 below shows the difference between the estimated and true repetitiveness as a function of the true repetitiveness, for ploidies 3, 4, 5, and 6.
- For genome length, we evaluated progenitor genomes of size 1 Mbp, 10 Mbp, 100 Mbp and 1 Gbp. We sequenced 15x coverage per homologue, and 1.0% sequencing error. Figure 6 below shows the relative error in the lengthas 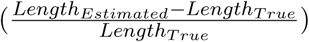 as a function of the true length (log scale), for ploidies 3, 4, 5 and 6.

**Figure 4:**
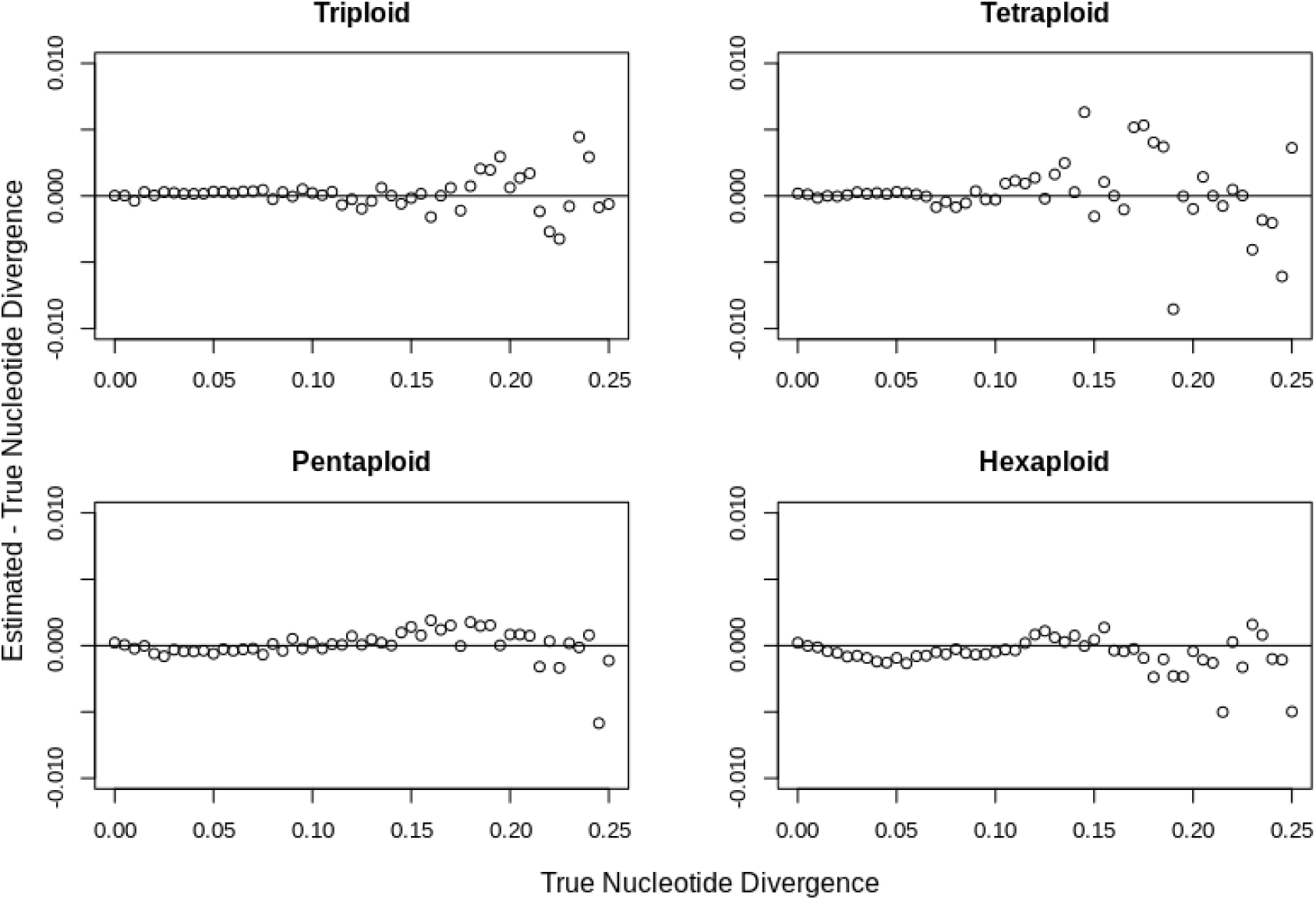
Nucleotide divergence parameter sweep for triploid, tetraploid, pentaploid, and hexaploid simulated datasets.

**Figure 5:**
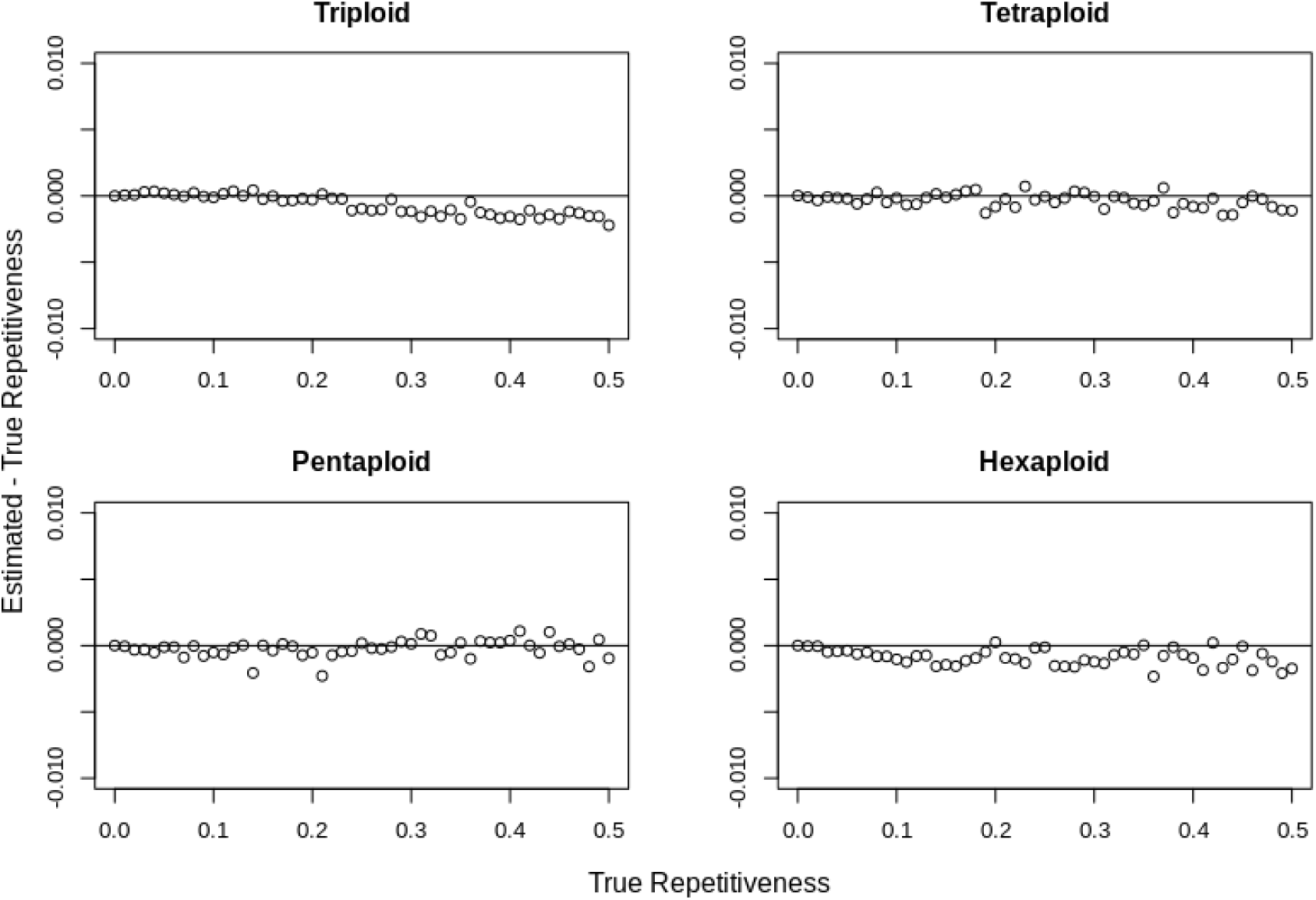
Repetitiveness parameter sweep for triploid, tetraploid, pentaploid, and hexaploid simulated datasets.

**Figure 6:**
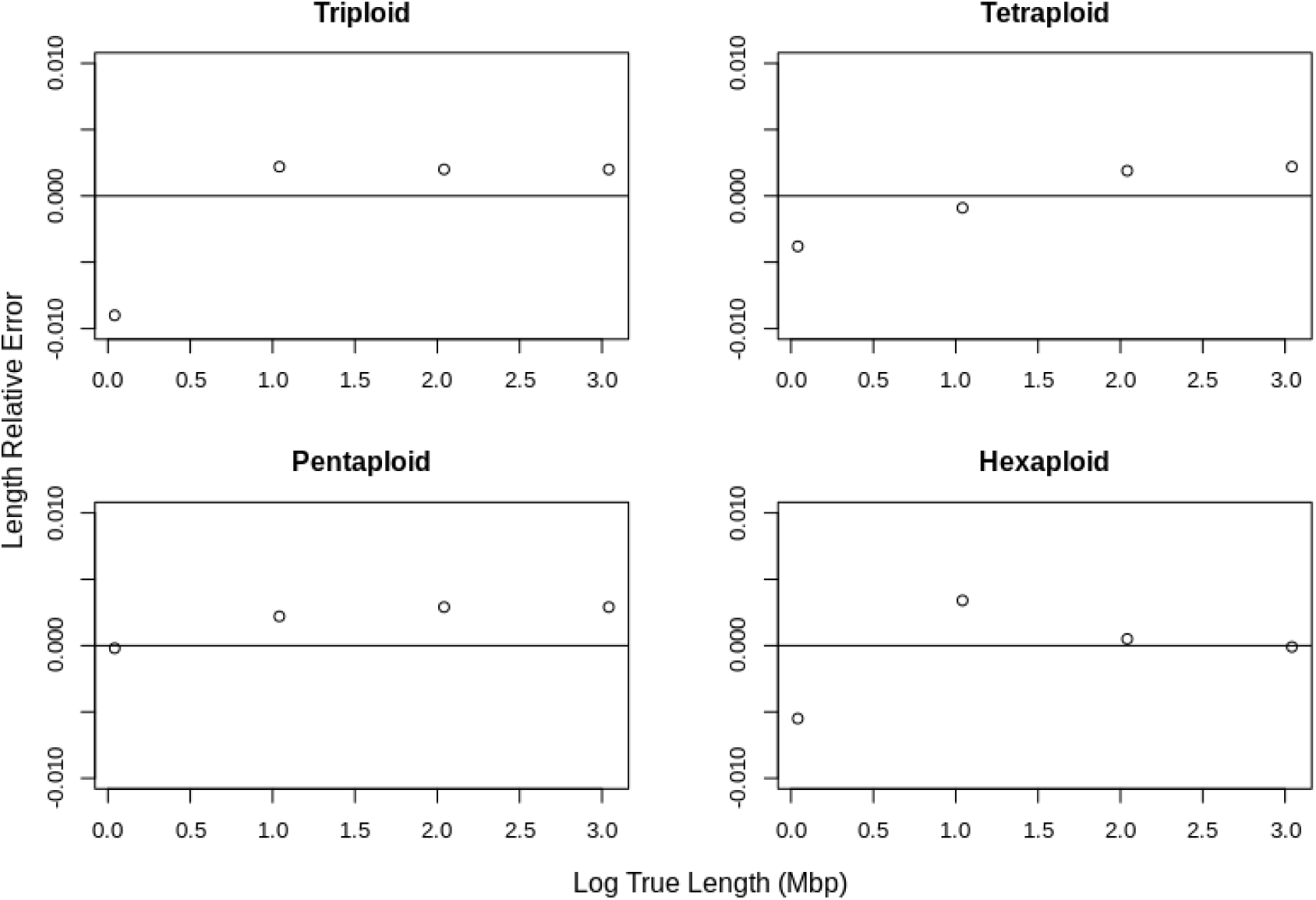
Length parameter sweep for triploid, tetraploid, pentaploid, and hexaploid simulated datasets.

### 5.3 Analysis of Real Polyploid Sequencing Data

We then applied GenomeScope 2.0 on the 11 real polyploid genomes listed in Table S1 (see Table 2 for the estimated polyploid genome sizes). Below we highlight a few notable results from this analysis, and the complete GenomeScope and Smudgeplot results are available within Section S3 in the **Online Methods**.

**Table 2:** Summary of polyploid genomes analyzed. The genome size refers to the polyploid genome size that is estimated by GenomeScope 2.0.

| Common Name | Species Name | Estimated Genome Size | Assembly Size |
| --- | --- | --- | --- |
| coastal redwood | *Sequoia sempervirens* | 26.9 Gbp | 26.5 Gbp |
| cotton | *Gossypium barbadense* | 2.291 Gbp | 2.267 Gbp |
| cotton | *Gossypium hirsutum* | 2.348 Gbp | 2.347 Gbp |
| marbled crayfish | *Procambarus virginalis* | 9.5 Gbp | 3.3 Gbp |
| root-knot nematode | *Meloidogyne arenaria* | 290.2 Mbp | 163.7 Mbp |
| root-knot nematode | *Meloidogyne enterolobii* | 268.6 Mbp | 162.4 Mbp |
| root-knot nematode | *Meloidogyne floridensis* | 201.7 Mbp | 74.9 Mbp |
| root-knot nematode | *Meloidogyne incognita* | 207.3 Mbp | 122.0 Mbp |
| root-knot nematode | *Meloidogyne javanica* | 280.1 Mbp | 142.6 Mbp |
| potato | *Solanum tuberosum* | 3.0 Gbp | 778.7 Mbp |
| wheat | *Triticum aestivum* | 14.1 Gbp | 15.34 Gbp |

Coastal redwoods (*Sequoia sempervirens*) are evergreen trees that can reach towering heights and are some of the longest living things on Earth. *Sequoia sempervirens* is known to be hexaploid, with recent evidence suggesting that it is an autohexaploid (Scott et al. 2016). This aligns with the Smudgeplot analysis, which inferred a triploid ploidy for this data, which comes from the haploid megagametophyte extracted from a seed. Furthermore, the genome size of the coastal redwood is larger than the human genome, with a recent assembly spanning 26.5 Gbp (Save the Redwoods League 2019). The estimated genome size of the coastal redwood output by GenomeScope is 26.9 Gbp, revealing great concordance with the recent assembly (see Figure S6 and Figure S7).

**Figure 7:**
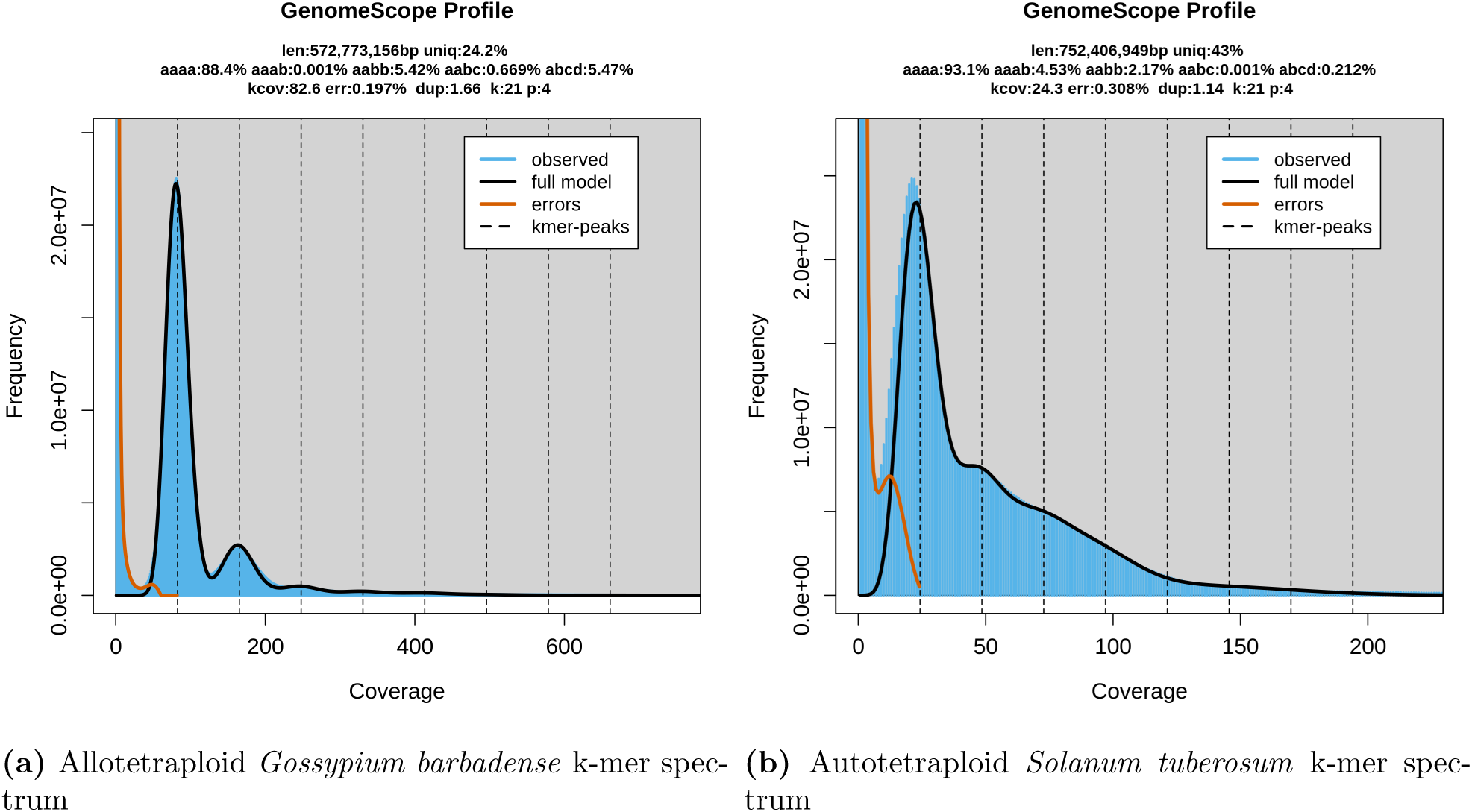
K-mer spectra for allotetraploid and autotetraploid species. Notice that the allotetraploid plot has *aaab < aaab*, while the autotetraploid plot has *aaab > aabb*.

Marbled crayfish (*Procambarus virginalis*) are freshwater crustaceans that undergo parthenogenetic reproduction, in which a female gamete develops into an individual without fertilization. Based on a Smudgeplot analysis, we inferred the ploidy to be triploid, which aligns with the current understanding of this organism (Gutekunst et al. 2018). We run GenomeScope 2.0 with a triploid model to estimate the genome characteristics. Specifically, GenomeScope estimates a polyploid genome size of 9.7 Gbp, while the current assembly spans 3.3 Gbp (see Figure S12 and Figure S13). It is clear that the assembly only spans one homologue of the triploid genome.

Root-knot nematodes (*Meloidogyne arenaria*, *Meloidogyne enterolobii*, *Meloidogyne floridensis*, *Meloidogyne incognita*, and *Meloidogyne javanica*) are parasitic roundworms that infect the roots of plants. Based on Smudgeplot analyses, we inferred that *Meloidogyne enterolobii*, *Meloidogyne floridensis*, and *Meloidogyne incognita* were triploid, while *Meloidogyne arenaria* and *Meloidogyne javanica* were tetraploid. Running GenomeScope 2.0 with the corresponding ploidies, we determined estimates for the genome characteristics. For the five root-knot nematodes the GenomeScope estimates for genome size are 1.65 to 2.69 times larger than the current best assemblies, suggesting the assemblies have partially collapsed the homologous chromosomes (see Figure S14 through Figure S23).

Bread wheat (*Triticum aestivum*) is an allohexaploid which consists of 3 subgenomes (IWGSC 2014). A Smudgeplot analysis inferred that the ploidy was diploid, because the individual subgenomes are highly divergent from each other. Specifically, if the homologous k-mers from different subgenomes are highly divergent (more than 1 SNP different), while the homologous k-mers from the same subgenome are only 1 SNP different, then we would expect to see three k-mer pairs. Each of these pairs would have an estimated sum of coverages of 2*λ* and an estimated relative minor coverage of ½, and would thus be interpreted by Smudgeplot as coming from the genomic structure *AB*. The current best assembly spans 15.34 Gbp, while the GenomeScope estimate is 14.1 Gbp (see Figure S26 and Figure S27).

### 5.4 Allotetraploid vs. Autotetraploid

One important application of GenomeScope is to distinguish between allotetraploid and autotetraploid species based on the distinct patterns of nucleotide heterozygosity rates that occur. For example, it is known in cotton that during meiosis homologous chromosomes from the same subgenome form bivalents and preferentially pair with each other (Endrizzi 1962). This phenomenon is also prominent in many other allotetraploid species (Xu et al. 2013). Thus, for allotetraploids we would expect a high proportion of *aabb* and a low proportion of *aaab* since preferential pairing would ensure that two homologues from the first subgenome and two homologues from the second subgenome are present after recombination. Conversely, it is known in potato that during meiosis the majority of cells contain quadrivalents (He et al. 2018). In this case, after recombination an individual might have 0, 1, 2 or 3 homologues from a given subgenome. Thus, *aaab* would be expected to be more prominent than *aabb* since it is more likely that there are 1 or 3 copies of a subgenome rather than exactly 2 copies of a subgenome.

For cotton and potato, we see that the GenomeScope estimates for nucleotide heterozygosity rates follow these expectations. For the two allotetraploid cotton species (Wang et al. 2019), *aaab* is estimated to be approximately 0 and *aabb* is estimated to be greater than 5%. The estimated genome size is also highly accurate, and GenomeScope estimates the polyploid genome length to be 2.291 Gbp and 2.348 Gbp while the current best assemblies span 2.267 Gbp and 2.347 Gbp respectively (see Figure S8 through Figure S11). For potato, *aaab* is greater than *aabb* as we would expect after recombination. Here the estimated genome size is approximately 3 times larger than the current best assembly (3.0Gbp vs 778.7Mbp) (see Figure S24 and Figure S25). This is expected since the assembly was filtered to form a pseudo-haploid representation that reports a single homolog (Hardigan et al. 2016). Thus, the GenomeScope estimates can determine whether a novel polyploid organism is an allopolyploid or autopolyploid.

## 6 Discussion

We have shown on simulated and real data sets that GenomeScope 2.0 is able to quickly and accurately estimate the genomic characteristics of polyploid organisms without a reference genome. The core of GenomeScope 2.0 is a polyploid model using the Möbius inversion formula which accounts for the k-mers occurring at higher ploidy levels. Users provide the k-mer spectrum as input, and GenomeScope performs a non-linear optimization using the Levenberg-Marquardt algorithm.

We have also introduced Smudgeplots as a visualization and analysis technique that can be used to reveal the structure of a novel species. The core of this analysis is the identification and statistical analysis of k-mer pairs that differ by exactly one nucleotide.

The coverage of the data set must be sufficient for these methods to resolve the error peak with the haploid peak. In general, having at least 15x coverage per homologue is sufficient. Relatedly, future work remains to extend these techniques for single molecule sequencing with high error rates that currently prevent k-mer based analysis. Species with both low heterozygosity and high repetitiveness may confuse a Smudgeplot analysis. For example, in the diploid *Fragaria iinumae* strawberry genome, more k-mer pairs come from the “AABB” smudge than from the “AB” smudge, which leads to the incorrect inference of tetraploidy (see Figure S24). Upon further analysis, Smudgeplot is correctly finding k-mer pairs in the genome, though they actually represent repetitive k-mer pairs, not k-mer pairs at a higher ploidy level. However, GenomeScope results reveal very low levels of heterozygosity and high rates of duplications, which highlights that using these tools in conjunction with one another can help unravel the properties of a genome.

Finally, polyploid species, especially allopolyploids, often have highly divergent genomic copies (e.g. *>* 12% different at the nucleotide level). Thus, one limitation of using a k-mer-based technique is that in these cases too few k-mers may actually be shared between the homologous copies. This can lead Smudgeplot to infer diploidy even for polyploid species. However, in these cases the divergence of the homologues may be so high that they will be separated during the assembly process. The polyploidy will then very likely be revealed by standard genome quality assessment of conserved single copy orthologs (BUSCO) (Simão et al. 2015).

Even with these caveats, GenomeScope and Smudgeplot are able to rapidly and accurately infer genomic properties for large, highly heterozygous, and polyploid genomes. GenomeScope accurately predicts genomic properties for the nearly 9 Gbp coastal redwood genome, for the highly heterozygous allotetraploid cotton genomes, and for the hexaploid wheat genome. Furthermore, GenomeScope is able to distinguish between allopolyploid and autopolyploid species, which can help researchers gain valuable biological insights for novel organisms without needing to perform costly experiments. Finally, Smudgeplot is able to correctly predict ploidy even in the extreme case of octaploid *Fragaria x ananassa*. These tools will open up future analysis of complex organisms that are underrepresented in current genomics pipelines.

## 7 Data availability

Genuine sequencing data are available using the accession codes listed in (Table S1). The code and parameters used for generating the simulated datasets is available in the GenomeScope 2.0 github repository. The full results of modeling the simulated datasets are available as a **Supplemental File**.

## 8 Code availability

All code supporting the current study is deposited in GitHub at https://github.com/tbenavi1/genomescope2.0 and https://github.com/KamilSJaron/smudgeplot. We also have an web-enabled version of GenomeScope available at http://genomescope.org/genomescope2.0/.

## Acknowledgements

We would like to thank Edward Scheinerman for his thorough explanation of combinatorics topics. We would also like to thank Tanja Schwander and Marc Robinson-Rechav for their helpful discussions. This work was supported, in part, by NIH grant R01-HG006677 and NSF grants DBI-1350041 and IOS-1732253 to MCS. KSJ was supported by Swiss National Foundation grant CRSII3 160723. Part of this research project was conducted using computational resources at the Maryland Advanced Research Computing Center (MARCC).

## 9 Author contributions

T.R.-B. extended the GenomeScope model for polyploid genomes. T.R.-B. and K.S.J. conceived and implemented Smudgeplots. M.C.S. supervised the project. T.R.-B., K.S.J., and M.C.S. wrote the manuscript.

## 11 Competing interests

The authors declare no competing interests.

## Online Methods

**Table S1:** Summary of polyploid genomes analyzed. The assembly size refers to the size of the assembly presented in the corresponding cited work.

| Common Name | Species Name | SRA | Ploidy | Assembly Size |
| --- | --- | --- | --- | --- |
| coastal redwood | *Sequoia sempervirens*<br>(Save the Redwoods League 2019) | SRR9087413<br>SRR9087414<br>SRR9087417<br>SRR9087419<br>SRR9087420<br>SRR9087425<br>SRR9087426<br>SRR9087428<br>SRR9087450<br>SRR9087484<br>SRR9087485<br>SRR9087486<br>SRR9087487<br>SRR9087508<br>SRR9087509<br>SRR9087510<br>SRR9087511<br>SRR9087512<br>SRR9087516<br>SRR9087517<br>SRR9087528<br>SRR9087529<br>SRR9087530<br>SRR9087531<br>SRR9087532<br>SRR9087533<br>SRR9087534<br>SRR9087535<br>SRR9087536<br>SRR9087537 | 6 | 26.5 Gbp |
| cotton | *Gossypium barbadense*<br>(Wang et al. 2019) | SRR1919013 | 4 | 2.267 Gbp |
| cotton | *Gossypium hirsutum*<br>(Wang et al. 2019) | SRX4734214 | 4 | 2.347 Gbp |
| marbled crayfish | *Procambarus virginalis*<br>(Gutekunst et al. 2018) | SRR5115143<br>SRR5115144<br>SRR5115145<br>SRR5115146<br>SRR5115147<br>SRR5115148 | 3 | 3.3 Gbp |
| root-knot nematode | *Meloidogyne arenaria*<br>(Szitenberg et al. 2017) | SRR4242457<br>SRR4242468<br>SRR4242476<br>SRR4242477 | 4 | 163.7 Mbp |
| root-knot nematode | *Meloidogyne enterolobii*<br>(Szitenberg et al. 2017) | SRR4242472<br>SRR4242473 | 3 | 162.4 Mbp |
| root-knot nematode | *Meloidogyne floridensis*<br>(Szitenberg et al. 2017) | SRR4242474<br>SRR4242475 | 3 | 74.9 Mbp |
| root-knot nematode | *Meloidogyne incognita*<br>(Szitenberg et al. 2017) | SRR4242460<br>SRR4242461 | 3 | 122.0 Mbp |
| root-knot nematode | *Meloidogyne javanica*<br>(Szitenberg et al. 2017) | SRR4242458<br>SRR4242459 | 4 | 142.6 Mbp |
| potato | *Solanum tuberosum*<br>(Hardigan et al. 2016) | SRR5349579 | 4 | 778.7 Mbp |
| wheat | *Triticum aestivum*<br>(Zimin et al. 2017) | SRX2994097 | 6 | 15.34 Gbp |

**Table S2:** Summary of estimated genome characteristics for the 11 analyzed real data sets. Genome size refers to the polyploid genome size. Heterozygosity refers to the nucleotide divergence. Repetitiveness refers to the percentage of the monoploid genome that consists of repetitive sequence.

| Species Name | Genome Size | Heterozygosity | Repetitiveness |
| --- | --- | --- | --- |
| *Sequoia sempervirens* | 26.9 Gbp | 4.4% | 53.5% |
| *Gossypium barbadense* | 2.291 Gbp | 11.6% | 75.8% |
| *Gossypium hirsutum* | 2.348 Gbp | 11.5% | 74.6% |
| *Procambarus virginalis* | 9.5 Gbp | 2.3% | 81.1% |
| *Meloidogyne arenaria* | 290.2 Mbp | 8.0% | 36.2% |
| *Meloidogyne enterolobii* | 268.6 Mbp | 6.1% | 38.1% |
| *Meloidogyne floridensis* | 201.7 Mbp | 2.8% | 24.4% |
| *Meloidogyne incognita* | 207.3 Mbp | 6.4% | 29.2% |
| *Meloidogyne javanica* | 280.1 Mbp | 8.4% | 35.2% |
| *Solanum tuberosum* | 3.0 Gbp | 6.9% | 57.0% |
| *Triticum aestivum* | 14.1 Gbp | 10.1% | 92.0% |

## S1 Combinatorial Model

### S1.1 Partially Ordered Sets

A partially ordered set, or *poset*, consists of a set *X* together with a binary relation *≤* satisfying reflexivity, anti-symmetry, and transitivity. Reflexivity states that for all *x ∈ X*, *x ≤ x*. Anti-symmetry states that for all *x, y ∈ X*, *x ≤ y* and *y ≤ x* implies *x* = *y*. Transitivity states that for all *x, y, z ∈ X*, *x ≤ y* and *y ≤ z* implies *x ≤ z*. A poset can be visualized by a directed acyclic graph in which the elements of the set are nodes in the graph and a directed edge exists from *x* to *y* if *x ≤ y*. To simplify this graph, it is common practice to depict only the direct edges and to ignore edges that can be implied by the transitive property.

Common examples of a poset include the real numbers with the standard less-than-or-equal relation, the integers with the divisibility relation, and the powerset of a set with the inclusion relation. An example of a poset with the inclusion relation is shown in Figure S1.

**Figure S1:**
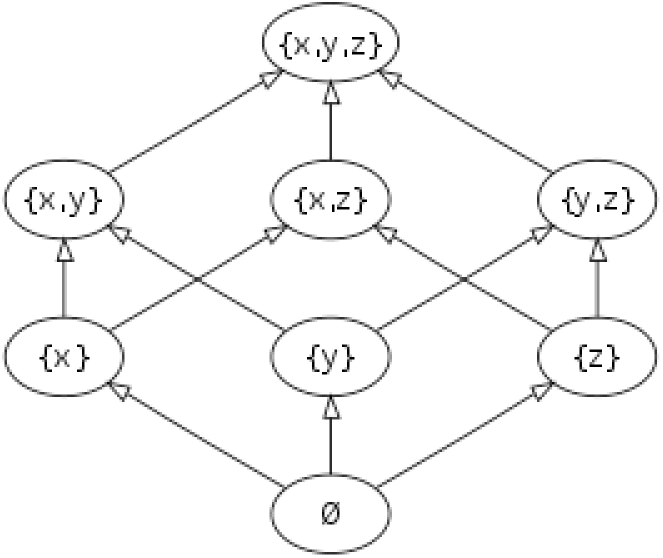
Inclusion poset on the set {*x, y, z*}.

### S1.2 Integer Partitions

For this problem, we apply a poset on integer partitions with the refinement relation. An integer partition of a positive integer *n* is a unordered tuple of positive integers such that their sum equals *n*. For example, (3, 1, 1, 1) is an integer partition of 6. We let Φ(*n*) denote the set of all integer partitions of *n*. We say that an integer partition *ϕ* is a refinement of the integer partition *ϕ′* if *ϕ* can be obtained by further partitioning elements of *ϕ′*, and we denote this by *ϕ ≤ ϕ′*. For example, (1, 1, 1, 1, 1, 1) *≤* (3, 1, 1, 1) because the element 3 can be partitioned into (1, 1, 1). The poset of the integer partitions of 4 is shown in Figure S2.

**Figure S2:**
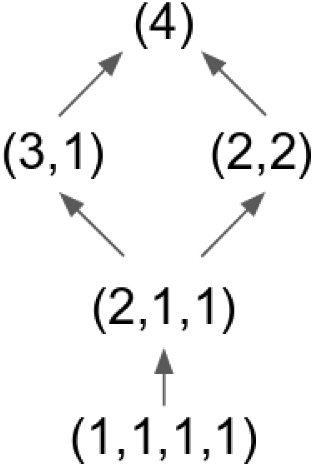
Poset of the integer partitions of 4.

### S1.3 Möbius Inversion Formula on Integer Partitions

Let *s* : Φ(*n*) *→* ℝ and *t* : Φ(*n*) *→* ℝ be real-valued functions defined on the integer partitions of *n*, with the property that 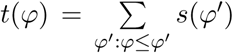. Furthermore, assume that calculating *t*(*ϕ*) is straightforward, but that we are actually interested in calculating *s*(*ϕ*). The Möbius inversion formula allows us to invert the above equation to calculate *s*(*ϕ*) in terms of *t*(*ϕ*):

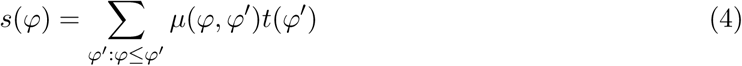

where *µ* is the Möbius function. The Möbius function is defined as

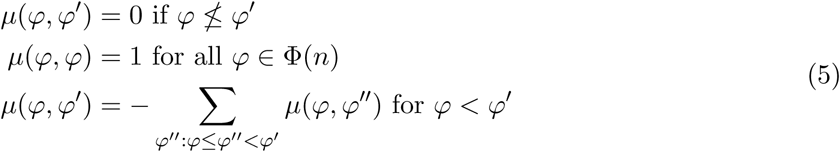

One useful property of Möbius functions is that they are defined based entirely on the poset structure, and are completely independent of the functions *s* and *t*.

### S1.4 Nucleotide Partitions

Recall the GenomeScope 2.0 polyploid model:

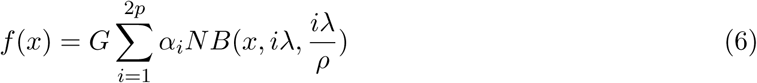

Now that we have introduced the necessary combinatorics theory, we more explicitly define the problem of determining *α_i_* in terms of the ploidy, repetitiveness, heterozygosity, and k-mer length. Let the ploidy *p* be the number of sets of homologous chromosomes and *x* be the number of chromosomes in a single complete set. We assume that, for each of the *x* chromosomes, all of the *p* corresponding homologues have exactly the same length.

**Figure S3:**
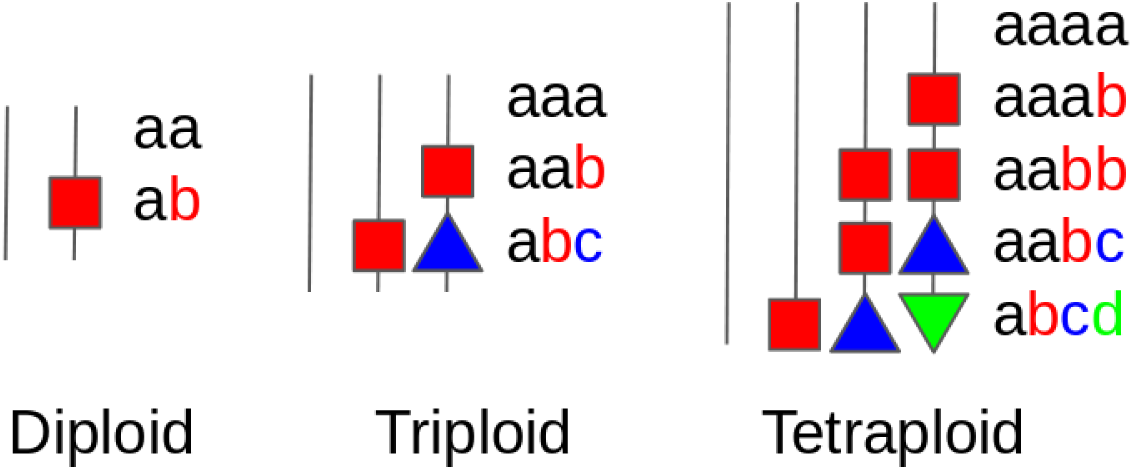
Nucleotide heterozygosity forms for the diploid, triploid, and tetraploid cases. The black vertical lines refer to the homologous chromosomes. The colored shapes correspond to distinct mutations that have accumulated on the homologues.

For any given position along the genome, the *p* nucleotides at that position may be homozygous or heterozygous (see Figure S3). In the diploid case, this corresponds to the nucleotides being all the same, *aa*, or the nucleotides being all different, *ab*. These correspond to the integer partitions (2) and (1, 1) respectively. In the polyploid case, however, there are more complicated possibilities. For example, in the triploid case it is possible for two nucleotides to be the same and the third to be different, *aab*, corresponding to the integer partition (2, 1).

In general, the nucleotides may group according to any of the integer partitions of *p*. Recall that the order of a nucleotide partition doesn’t matter, so *aba* and *aab* are equivalent. Indeed, this makes sense for our problem since the data in a k-mer spectrum are not homolog-specific and it is mathematically impossible to distinguish between equivalent nucleotide partitions.

### S1.5 Nucleotide Heterozygosity Rates

For our model, we assume the infinite sites model. Thus, we can define nucleotide heterozygosity rates corresponding to the probabilities that the nucleotides across the *p* homologues at a given location of the genome partition according to a given integer partition. We define *r_ϕ_* as the nucleotide heterozygosity rate corresponding to the nucleotide partition *ϕ*. For example, in the diploid case, the nucleotide heterozygosity rate, *r*_(1,1)_, corresponds to the probability that the two nucleotides at a given position in the genome are distinct, i.e. that they partition according to *ab*. The nucleotide homozygosity rate, *r*_(2)_, corresponds to the probability that the two nucleotides partition according to *aa* and is given by *r*_(2)_ = 1 *− r*_(1,1)_.

Similarly, in the polyploid case, the nucleotide heterozygosity rates are defined according to the nucleotide partitions. For example, in the hexaploid case, *r*_(3,2,1)_ corresponds to the probability that the nucleotides partition according to *aaabbc*. The nucleotide homozygosity rate, *r*_(6)_, corresponds to the probability that the nucleotides partition according to *aaaaaa*, and is given by 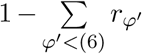.

### S1.6 K-mer Partitions

As the k-mer spectrum deals with k-mers and not with individual nucleotides, it is necessary to relate nucleotide heterozygosity rates with k-mer partition rates. Let *k* correspond to the k-mer length. Note that for any position along the genome (except for the final *k −* 1 positions on each chromosome), the *p* k-mers beginning at this position may group according to any of the integer partitions of *p*. Similar to nucleotide partitions, the order of k-mer partitions doesn’t matter, so *ABA* is equivalent to *AAB*. Furthermore, as with nucleotide partitions, it is mathematically impossible to distinguish between equivalent k-mer partitions in the k-mer spectrum.

### S1.7 K-mer Heterozygosity Rates

We define k-mer heterozygosity rates corresponding to the probabilities that the k-mers across the *p* homologues at a given location of the genome partition according to a given integer partition. We define *s_ϕ_* as the k-mer nucleotide heterozygosity rate corresponding the the k-mer partition *ϕ*. In the diploid case, the k-mer partition rates *s*_(2)_ and *s*_(1,1)_ correspond to the probabilities that the two k-mers at a given position (in a non-repetitive region of the genome) partition according to *AA* and *AB* respectively. Note that the only way for the k-mers to partition according to *AA* is if, for each of the *k* positions along the k-mer, the nucleotides partition according to *aa* (see Figure S4). Thus, assuming the infinite sites model, *s*_(2)_ = (*r*_(2)_)*^k^*, which is equivalent to the more general form:

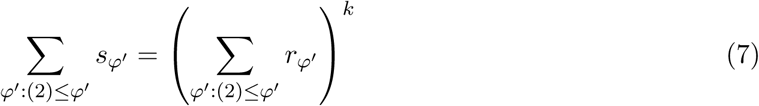

**Figure S4:**
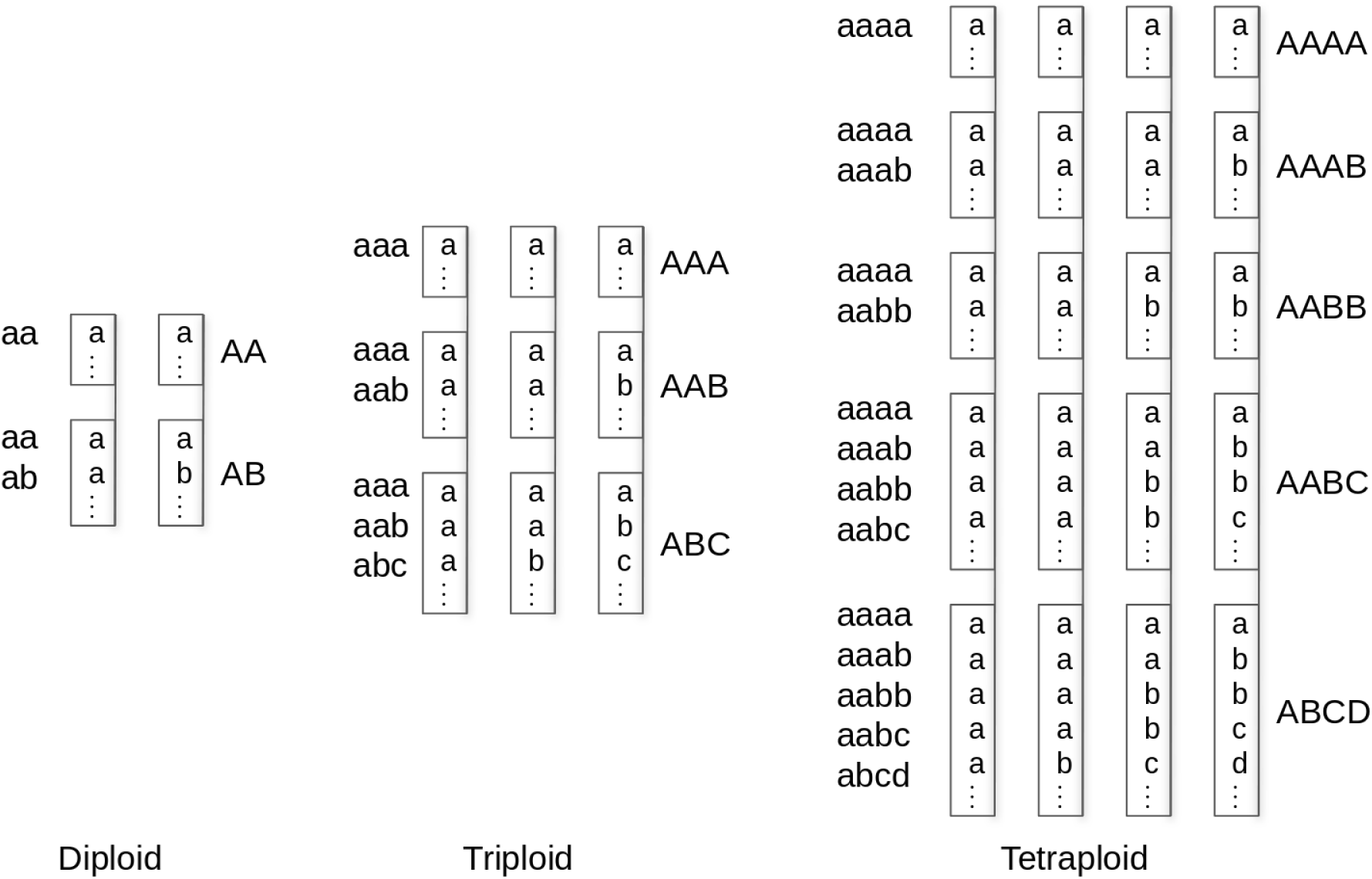
K-mer heterozygosity forms and their corresponding nucleotide heterozygosity forms in the diploid, triploid, and tetraploid cases. The black vertical lines refer to the homologous chromosomes. The black boxes refer to the k-mers on the homologues. The nucleotide heterozygosity forms on the left are compatible with the k-mer heterozygosity form on the right. Specifically, the k-mers will partition according to the k-mer partition on the right, as long as they are made up of any combination of nucleotides partitioned according to the nucleotide heterozygosity forms on the left.

To determine *s*_(1,1)_, one must consider which nucleotide partitions are compatible with the k-mer partition *AB*. In fact, both *ab* and *aa* are compatible. For example, consider the k-mers *gattaca* and *cattaca*. These k-mers are distinct and thus partition according to *AB*. However, while the nucleotides at the first position partition according to *ab*, the nucleotides at positions two through seven partition according to *aa*. Thus, (*r*_(1,1)_ + *r*_(2)_)*^k^*, which represents the probability that the nucleotides at every position along the k-mer partition according to *ab* or *aa*, is equivalent to the probability that the k-mers partition according to *AB* or *AA*. This yields

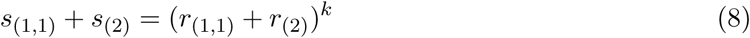

which is equivalent to the more general form

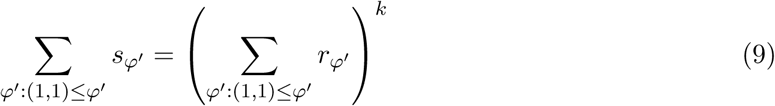

This further implies

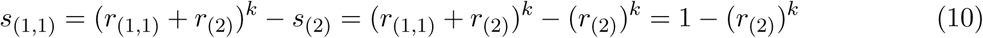

In the general polyploid case, it is possible to determine which nucleotide partitions are compatible with a given k-mer partition by using the integer partition poset. Specifically, any nucleotide partition *ϕ* in the poset is compatible with any k-mer partition *ϕ′* in the poset if and only if *ϕ ≥ ϕ′*. For example, returning to *gattaca* and *cattaca*, we have that *aa* is compatible with *AB* since (2) ≥ (1, 1).

Let 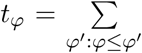 represent the probability that the k-mers partition according to *ϕ* or any other partition *ϕ* with *ϕ < ϕ′*. This is straightforward to calculate in terms of nucleotide partition rates as 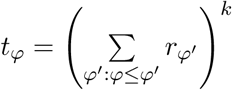.

### S1.8 Applying the Möbius Inversion Formula

Using the Möbius inversion formula, we can calculate *s_ϕ_*in terms of *t_ϕ_*. Specifically, we have

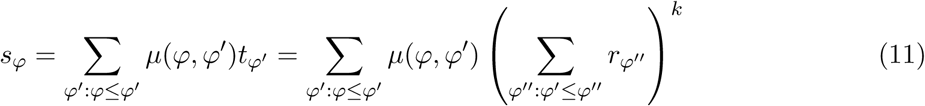

which gives us the k-mer heterozygosity rates in terms of the nucleotide heterozygosity rates.

### S1.9 K-mer Frequency Contributions in Non-Repetitive Regions

With these equations derived for the k-mer partition rates, it is necessary to determine how the *p* k-mers of each of the possible k-mer partitions contribute to the 2*p* peaks of the k-mer spectrum. Let *M_i_*(*ϕ*) denote the frequency contribution to peak *i* by the *p* k-mers (in a non-repetitive region) partitioned according to *ϕ*. For example, if *ϕ* = *AAABBCCD*, then *M*_1_(*ϕ*) = 1 because the *D* k-mer contributes to the first peak, *M*_2_(*ϕ*) = 2 since the *B* and *C* k-mers contribute to the second peak, and *M*_3_(*ϕ*) = 1 since the *A* k-mer contributes to the third peak.

### S1.10 K-mer Frequency Contributions in Repetitive Regions

For k-mers that are a two-copy repeat, there are two locations of the genome where they occur. Let *ϕ*_1_ be the k-mer partition of the *p* k-mers at the first location, and *ϕ*_2_ be the k-mer partition of the *p* k-mers at the second location. We make the simplifying assumption that the repetitive k-mer (i.e. the k-mer that is equivalent between the two k-mer partitions) is the most prevalent k-mer in each of the two k-mer partitions. For example, if *ϕ*_1_ = *AAABBC* and *ϕ*_2_ = *AABBCC*, then the overall k-mer partition of the 2*p* k-mers is *AAAAABBCCDDE*. Specifically, we consider the *A* k-mers between partitions to be equivalent, but not the *B* and *C* k-mers. Then, we may let *M_i_*(*ϕ*_1_*, ϕ*_2)_ denote the frequency contribution to peak *i* by the 2*p* k-mers (in a two-copy repeat) partitioned according to *ϕ*_1_ and *ϕ*_2_.

### S1.11 Polyploid Alpha Coefficients

Finally, we have:

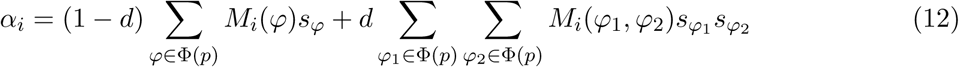

where *d* is the proportion of distinct k-mers of the monoploid genome that occur twice, *p* is the ploidy, Φ(*p*) is the set of integer partitions of *p*, *M_i_*(*ϕ*) and *M_i_*(*ϕ*_1_*, ϕ*_2)_ are the frequency contributions to peak *i* of the k-mers partitioned according to *ϕ* or (*ϕ*_1_*, ϕ*_2)_ respectively, and *s_ϕ_* is the k-mer heterozygosity rate of the k-mer partition *ϕ*.

## S2 Topologies

In the field of phylogenetics, the evolutionary relationships between species are often depicted in a branching diagram known as a phylogenetic tree. In this setting, the topology of the tree refers to the branching structure of the tree. We may also depict the similarities between homologous chromosomes in a branching diagram. In this case, a topology refers to the similarities between distinct homologues.

For ploidies of 4 and greater, there are multiple possible topologies (see Figure S5). For example, the two tetraploid topologies are *AAAA → AAAB → AABC → ABCD* and *AAAA → AABB → AABC → ABCD*.

**Figure S5:**
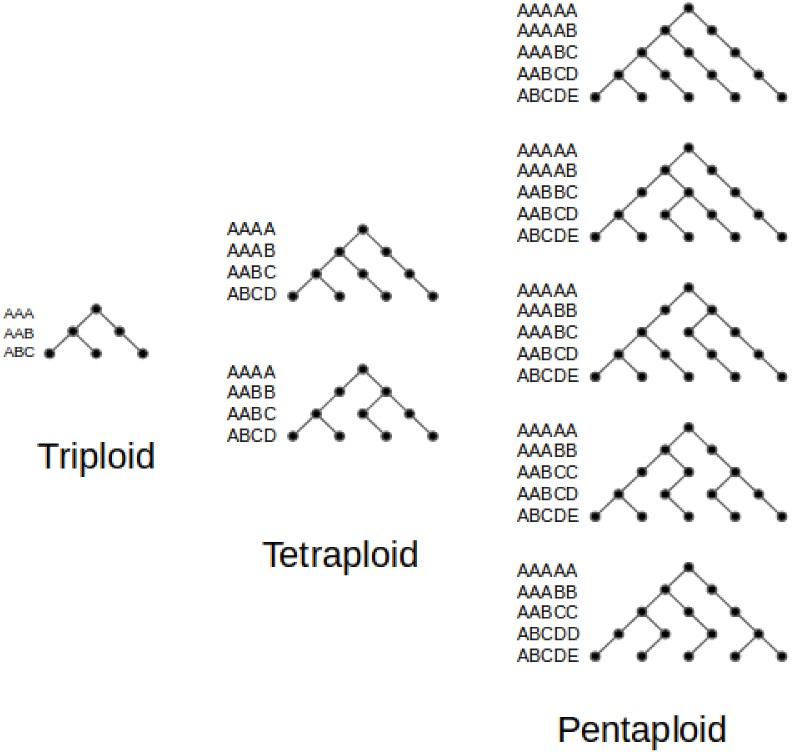
Topologies in the triploid, tetraploid, and pentaploid cases.

### S2.1 Tetraploid Topologies

For an autotetraploid organism, a whole genome duplication event has occurred sometime in its evolutionary history. Thus, for a given locus, the two k-mers at this locus of the ancestral genome were either heterozygous or homozygous (for an ancestral mutation) at the time of duplication. If the ancestral k-mers were homozygous at this locus, then the four k-mers of the polyploid organism immediately after the duplication were of the form AAAA.

Now we must consider the possibility that a more recent mutation that overlaps the k-mers at this locus has accumulated in the population. In this case, after recombination a sequenced individual may have this new mutation in zero, one, two, three, or four homologues. If this new mutation occurs in one or three homologues, then the k-mers are of the form AAAB. If this new mutation occurs in two homologues, then the k-mers are of the form AABB. Notably, AAAB is more prevalent than AABB because it is more likely that a mutation will be on any one homologue or any three homologues (4*p*(1 *− p*)^3^ + 4*p*^3^(1 *− p*)) versus any two homologues (6*p*^2^(1 *− p*)^2^), where *p* is the allele frequency of the mutation in the population.

If instead the ancestral k-mers were heterozygous at this locus (which is much rarer than the k-mers being homozygous at this locus), then the four k-mers of an ancient polyploid organism immediately after duplication were of the form AABB. For a modern organism which has undergone recombination, this ancestral mutation may be present in any number of the four homologues.

If the ancestral mutation is present in zero or all four homologues, then the k-mers (disregarding modern mutations) are of the form AAAA. Again, we must then consider that a more recent mutation may be present in any number of homologues of a sequenced individual. If the recent mutation is present in one or three homologues, then the k-mers are of the form AAAB, while if it is present in two homologoues, then the k-mers are of the form AABB. Again, AAAB would be more prevalent than AABB due to the same reasoning as above.

Finally, if the ancestral mutation were present in one or three homologues, then the k-mers were of the form AAAB, while if it were present in two homologues, then the k-mers were of the form AABB. Again, AAAB would be more prevalent than AABB. In summary, we would expect that the prevalence of AAAB would be much greater than the prevalence of AABB in autotetraploid species.

Intuitively, the only ways for the k-mers to partition according to AABB in an autotetraploid species are 1) the k-mers were homozygous before the duplication event and any modern mutations have accumulated on exactly two homologues after recombination or 2) the k-mers were heterozygous before the duplication event and the the ancient mutation has accumulated on exactly two homologues after recombination and any modern mutation has accumulated on the same two homologues or on the opposite two homologues. For this reason, we would expect that the k-mer heterozygosity rate of AABB in autotetraploid species lower than that of AAAB, and define the “autotetraploid topology” as *AAAA → AAAB → AABC → ABCD*.

For an allotetraploid organism, two similar but distinct ancestral species have undergone a hybridization event sometime in its evolutionary history. Thus, for a given locus, the two k-mers of the first ancestral genome may either be heterozygous or homozygous (for an ancestral mutation) and the two k-mers of the second ancestral genome may either be heterozygous or homozygous (for another ancestral mutation). If the k-mers at this locus in both ancestral genomes were homozygous, which is quite likely, then we would expect the k-mers to be of the form AABB. Furthermore, due to the preferential chromosomal pairing of A with A and B with B that is often the case during meiosis with allotetraploid species, we would still expect a high prevalence of AABB after recombination.

Thus in the allotetraploid case, AABB is more prevalent because it is much more likely that the k-mers at a particular locus in the ancestral genomes were homozygous rather than heterozygous and because it is much more likely that homologous chromosomes from the same ancestral species pair together during meiosis. Intuitively, the reason why AABB is more prevalent for allotetraploid species than for autotetraploid species is because for allotetraploid species there are two distinct genomes. Thus, homozygous locations of the genome can result in AABB, whereas for autotetraploid species there is only a single duplicate genome so homozygous locations necessarily result in AAAA. In this case AABB is then only possible for an autotetraploid species if a more recent mutation occurs in exactly two homologues. In summary, we would expect that the prevalence of AABB would be much greater than the prevalence of AAAB in allotetraploid species. For this reason, we define the “allotetraploid topology” as *AAAA → AABB → AABC → ABCD*.

## S3 Analysis of Polyploids

### S3.1 Coastal Redwood Results

**Figure S6:**
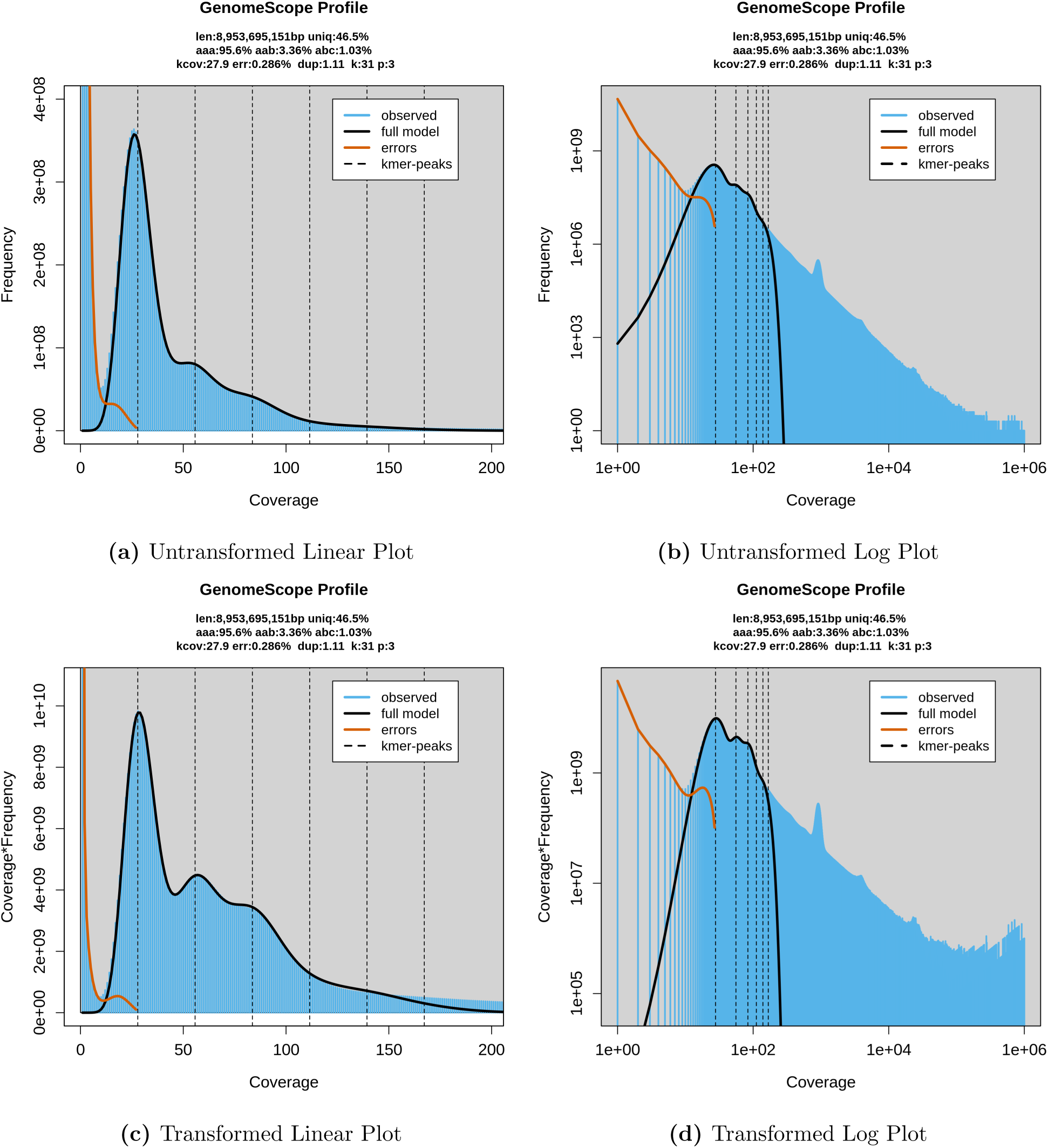
GenomeScope results for *Sequoia sempervirens*. Note that while the coastal redwood is hexaploid, these data are actually triploid since they come from the megagametophyte extracted from a seed.

**Figure S7:**
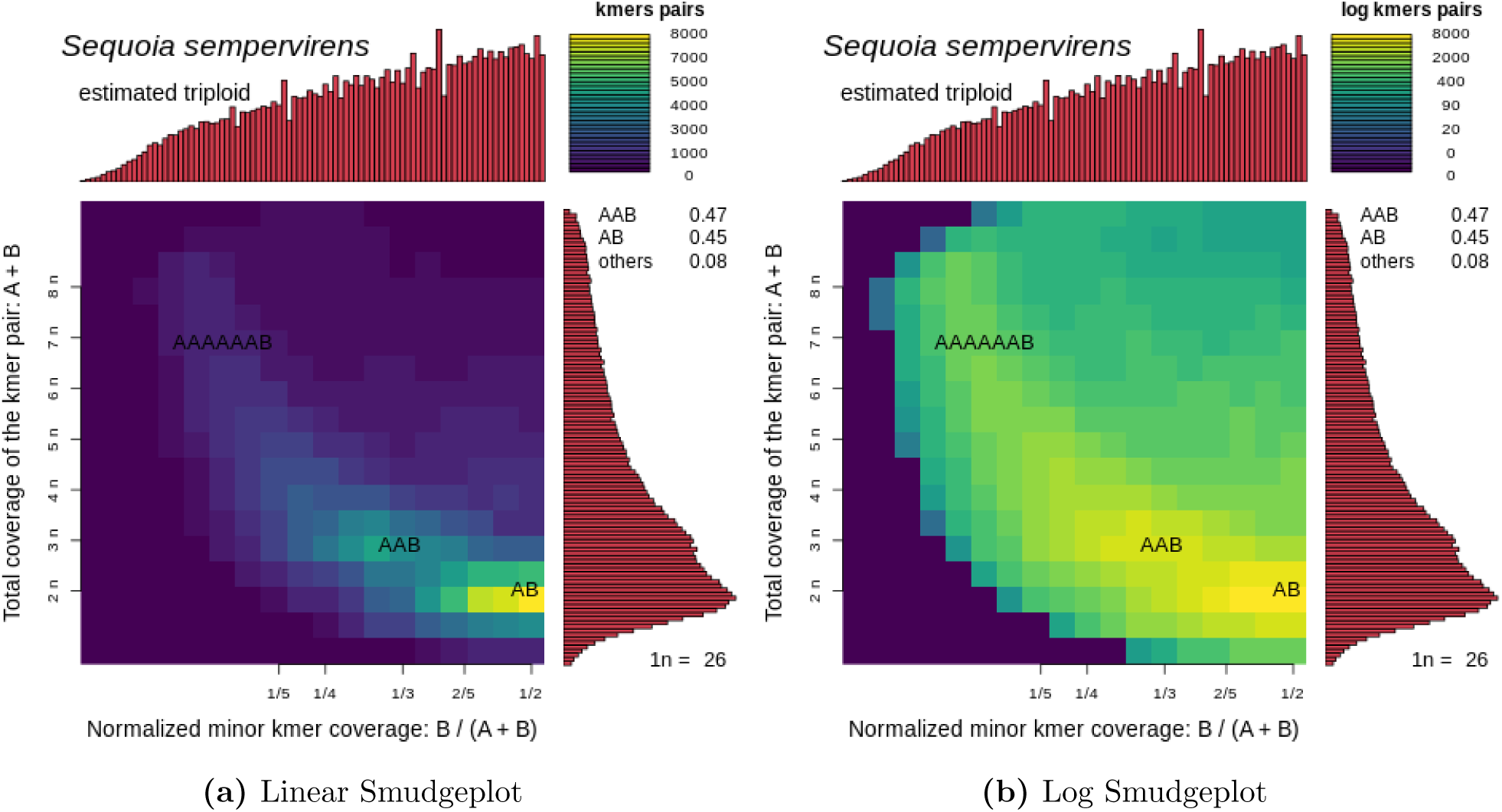
Smudgeplot results for *Sequoia sempervirens*.

### S3.2 Cotton Results

**Figure S8:**
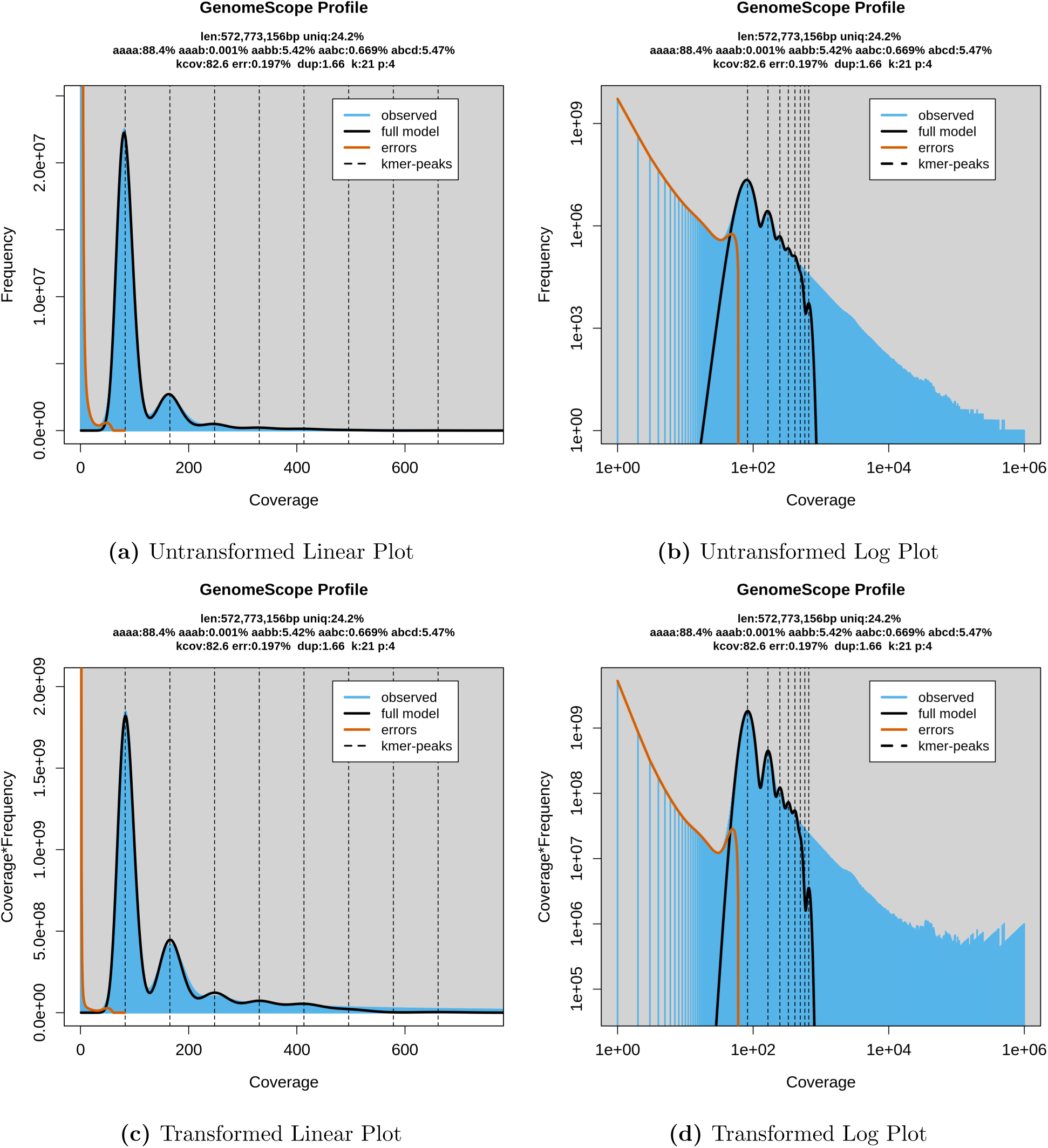
GenomeScope results for *Gossypium barbadense*.

**Figure S9:**
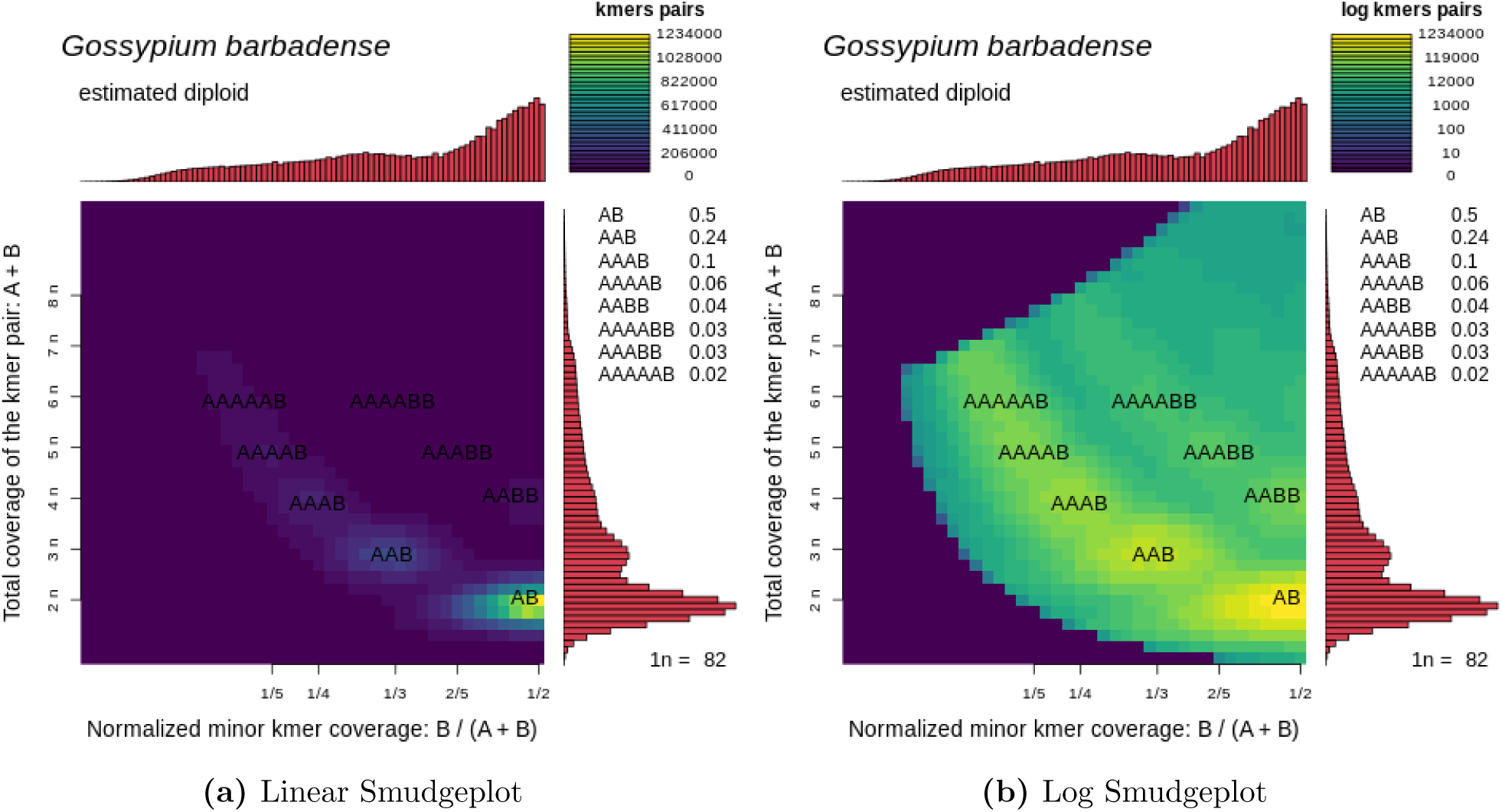
Smudgeplot results for *Gossypium barbadense*.

**Figure S10:**
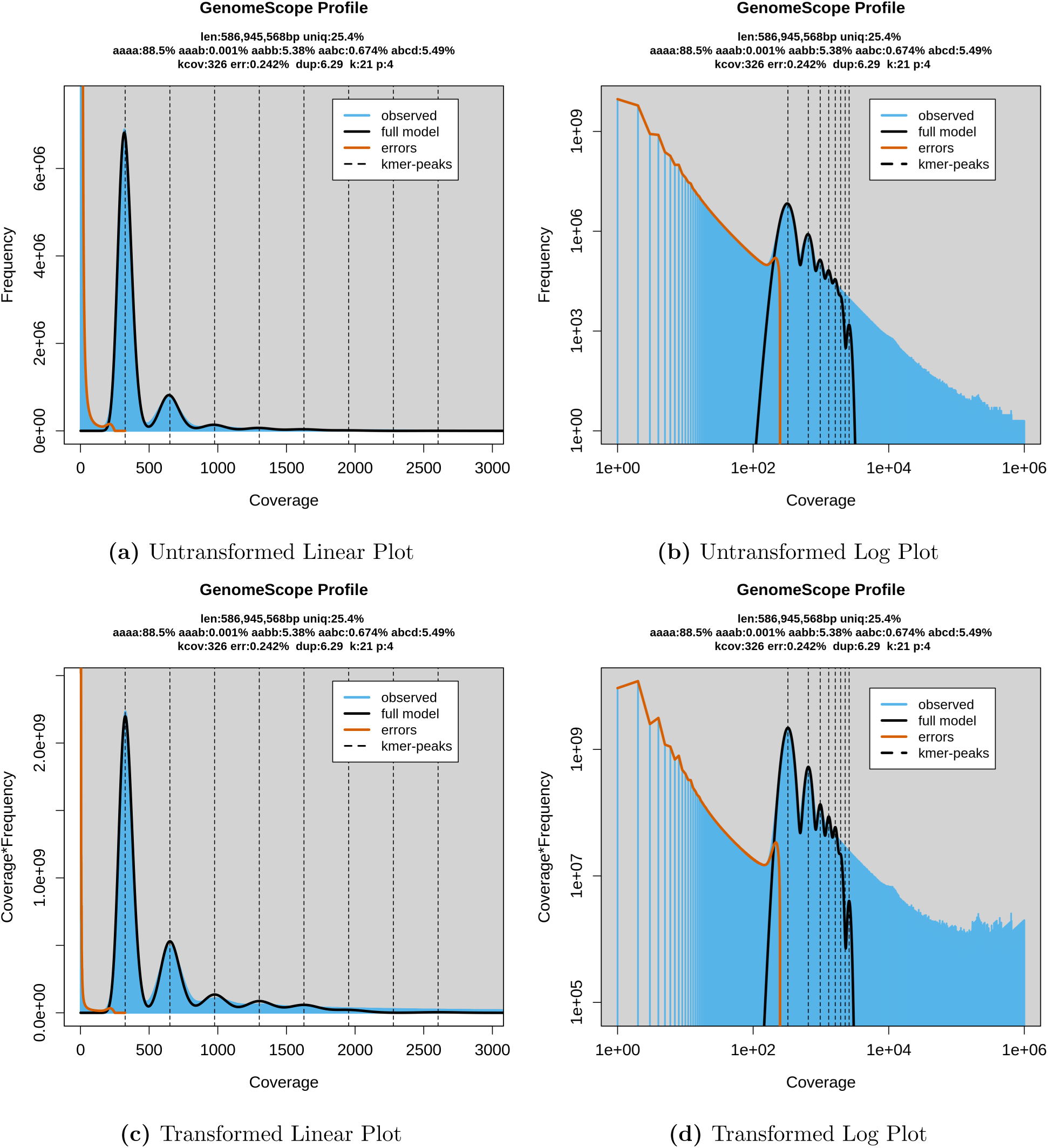
GenomeScope results for *Gossypium hirsutum*.

**Figure S11:**
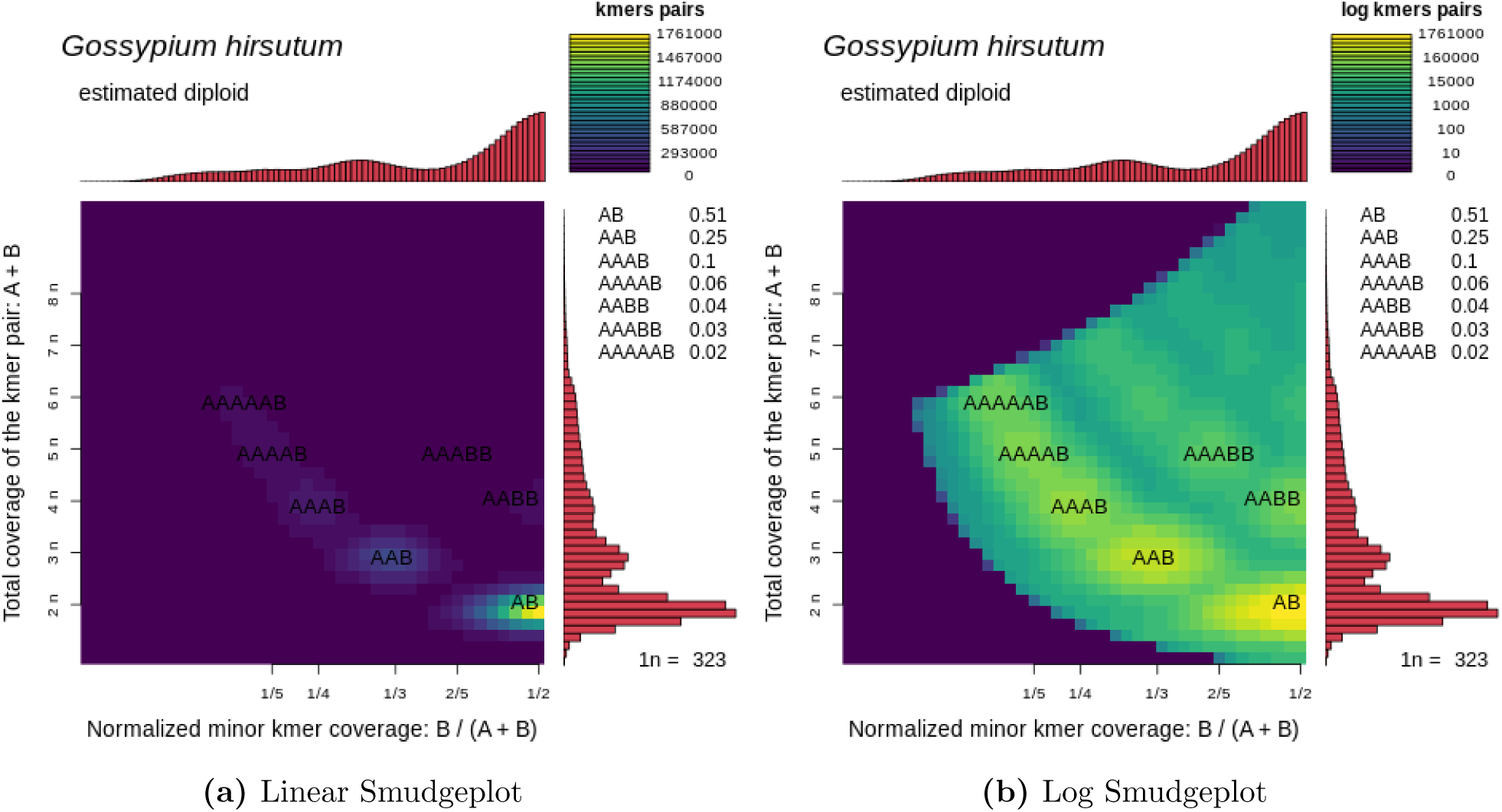
Smudgeplot results for *Gossypium hirsutum*.

### S3.3 Marbled Crayfish Results

**Figure S12:**
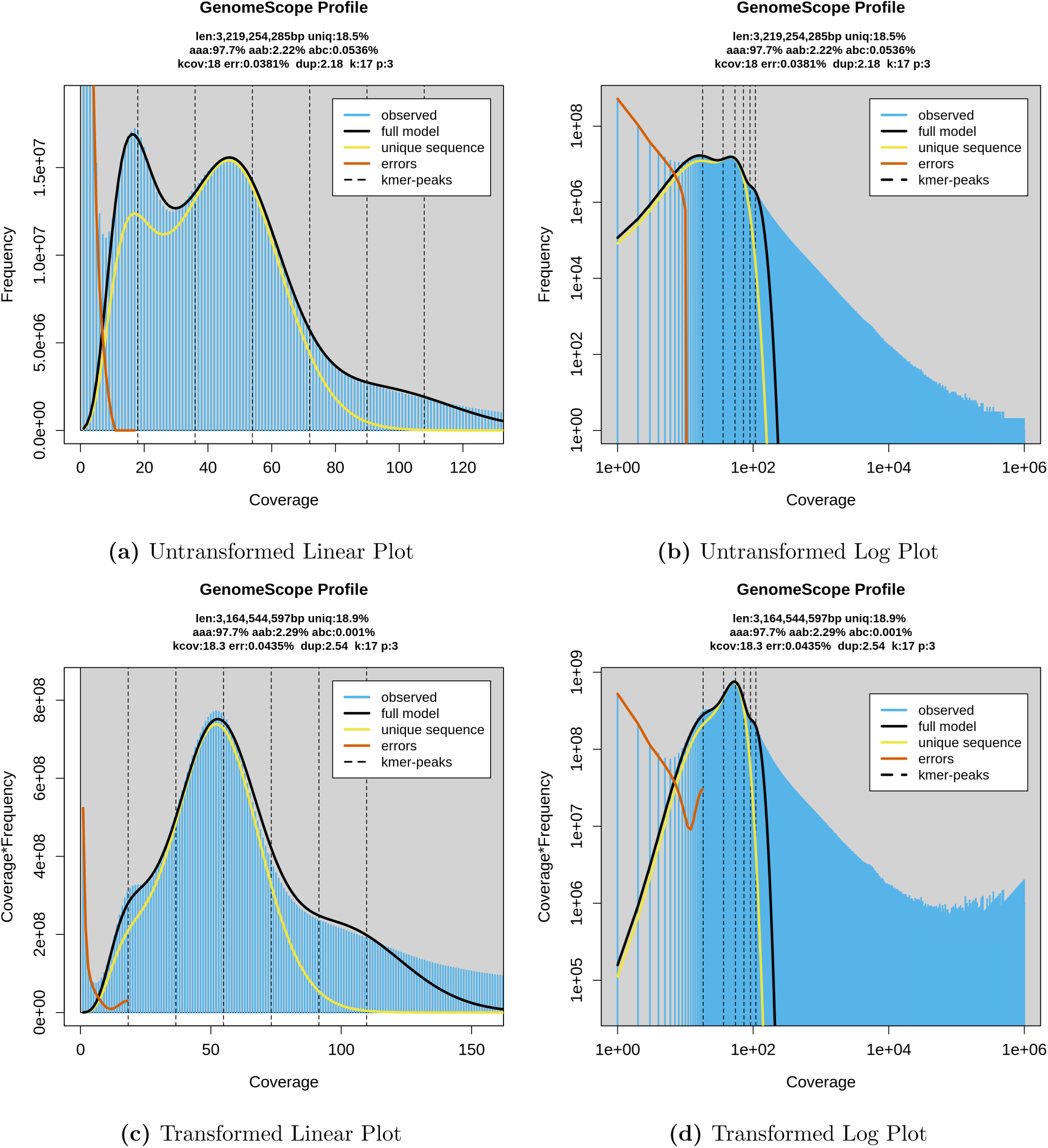
GenomeScope results for *Procambarus virginalis*.

**Figure S13:**
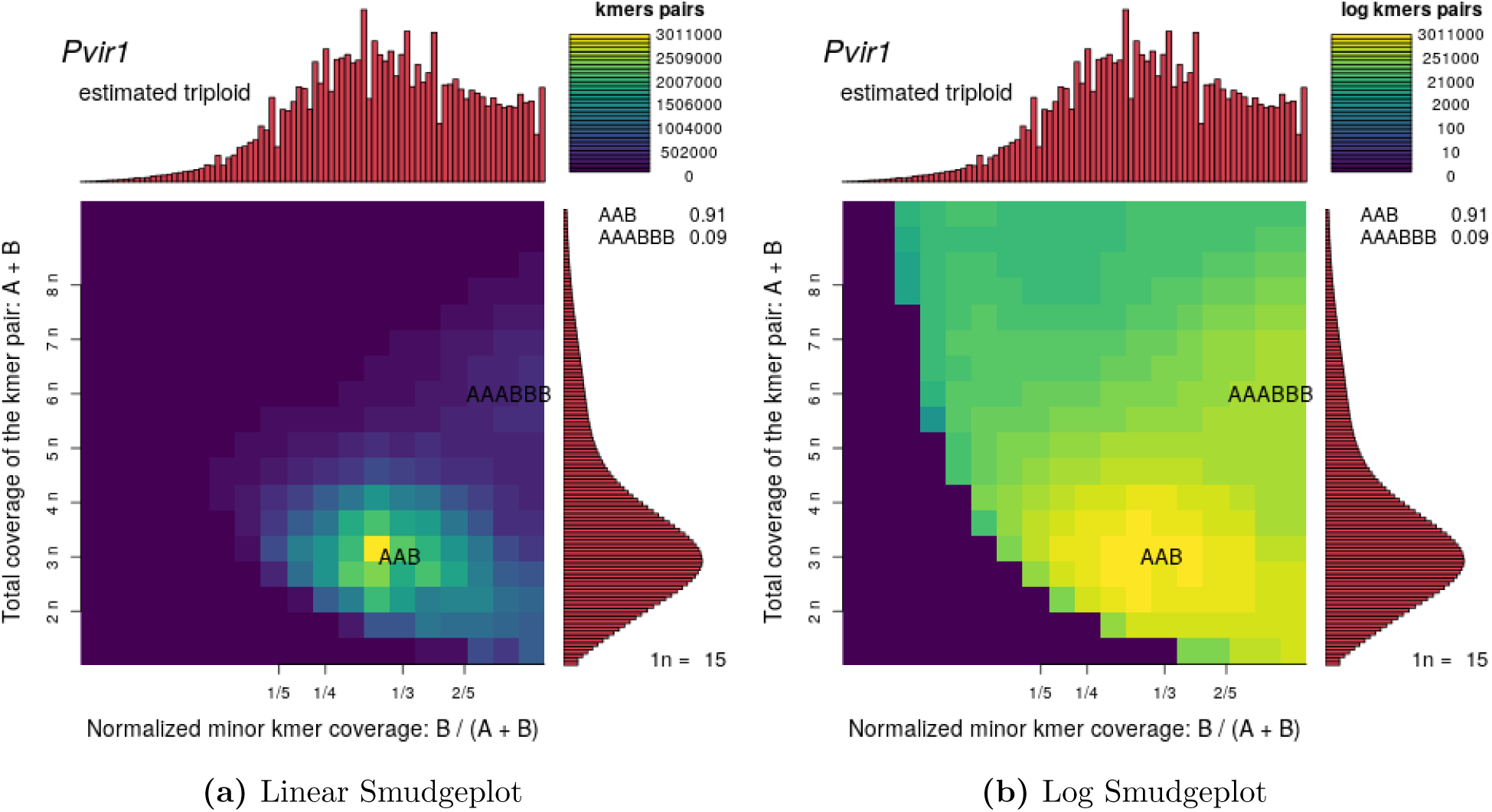
Smudgeplot results for *Procambarus virginalis*.

### S3.4 Root-knot Nematode Results

**Figure S14:**
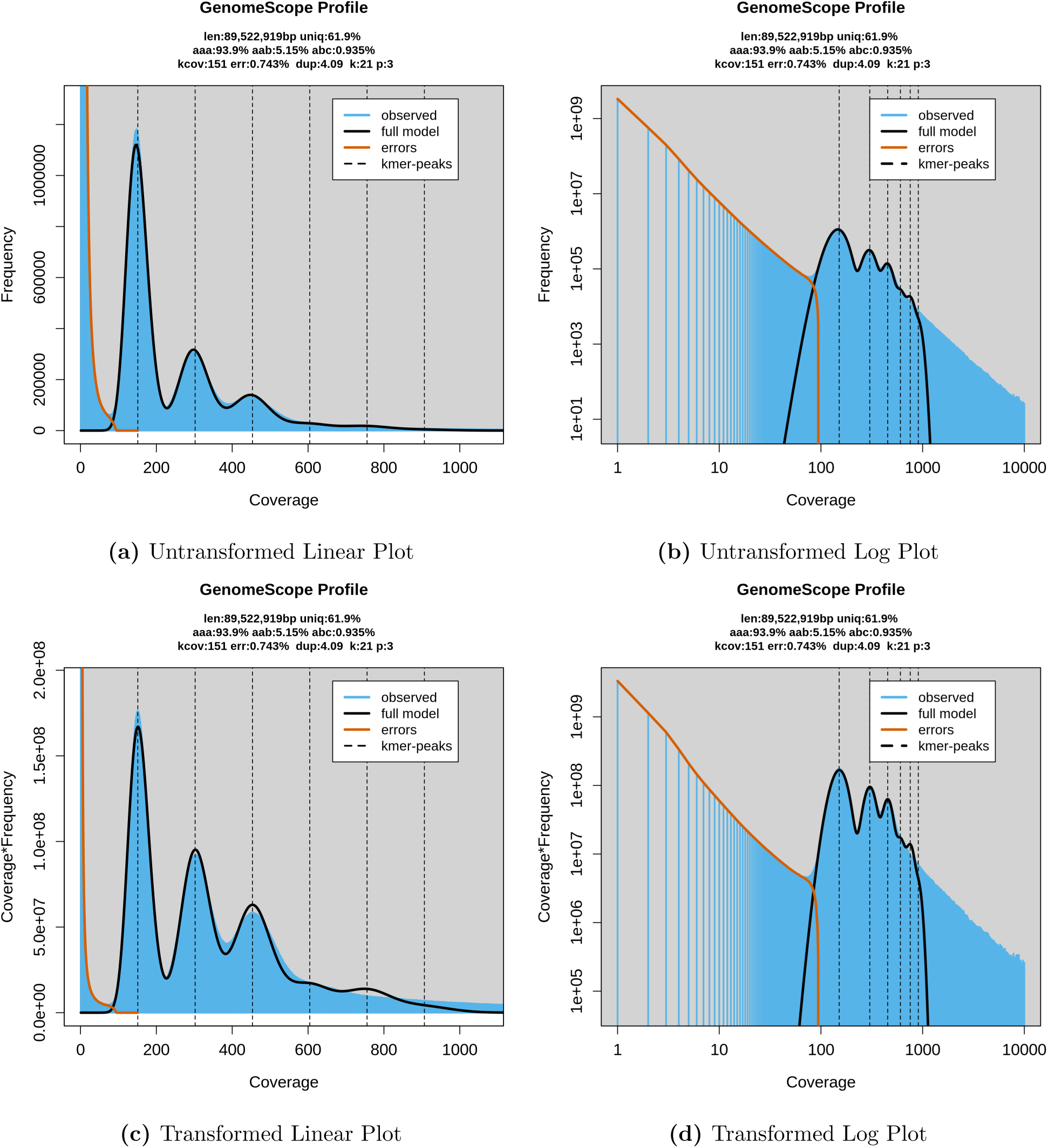
GenomeScope results for *Meloidogyne enterolobii*.

**Figure S15:**
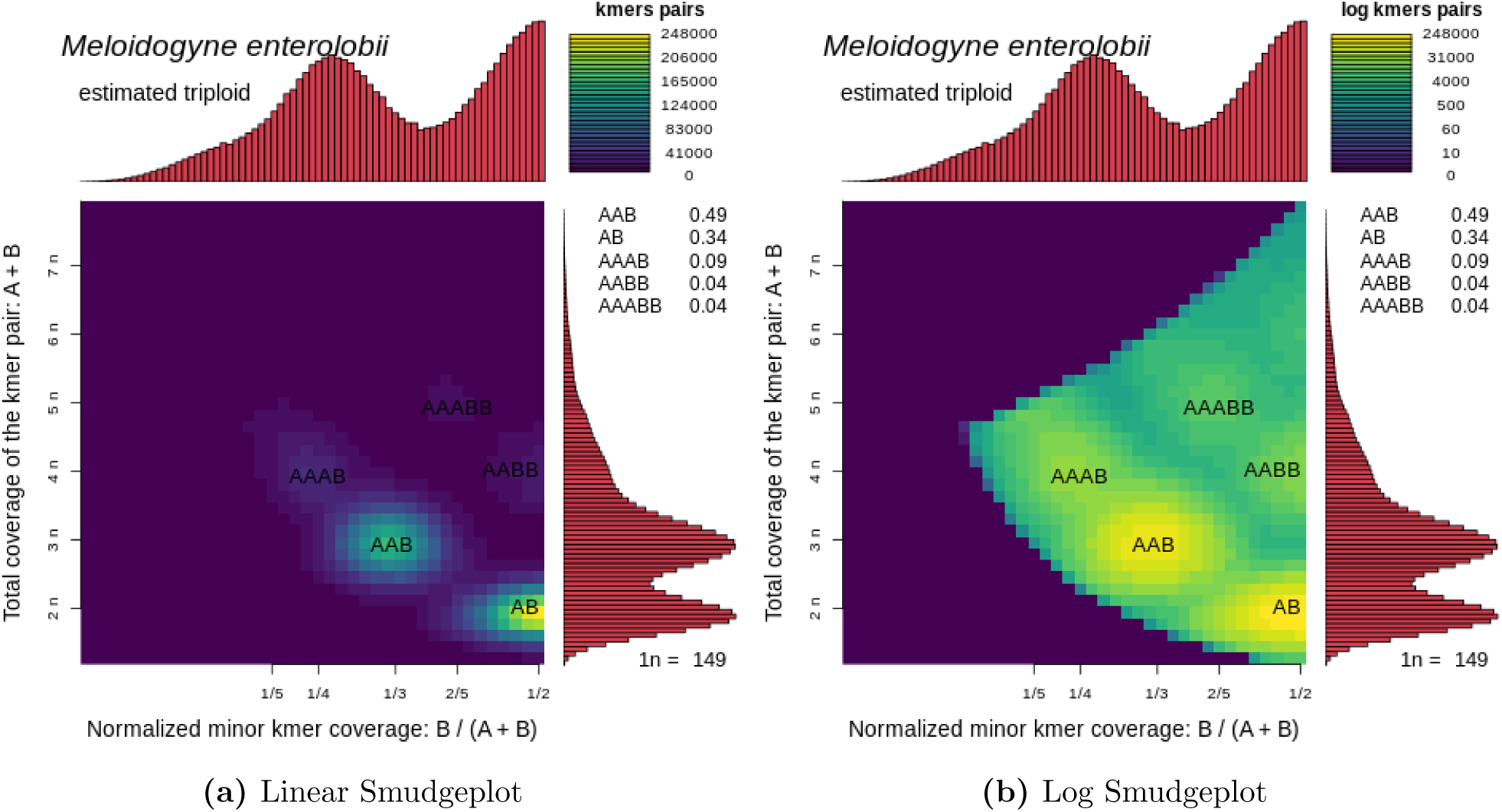
Smudgeplot results for *Meloidogyne enterolobii*.

**Figure S16:**
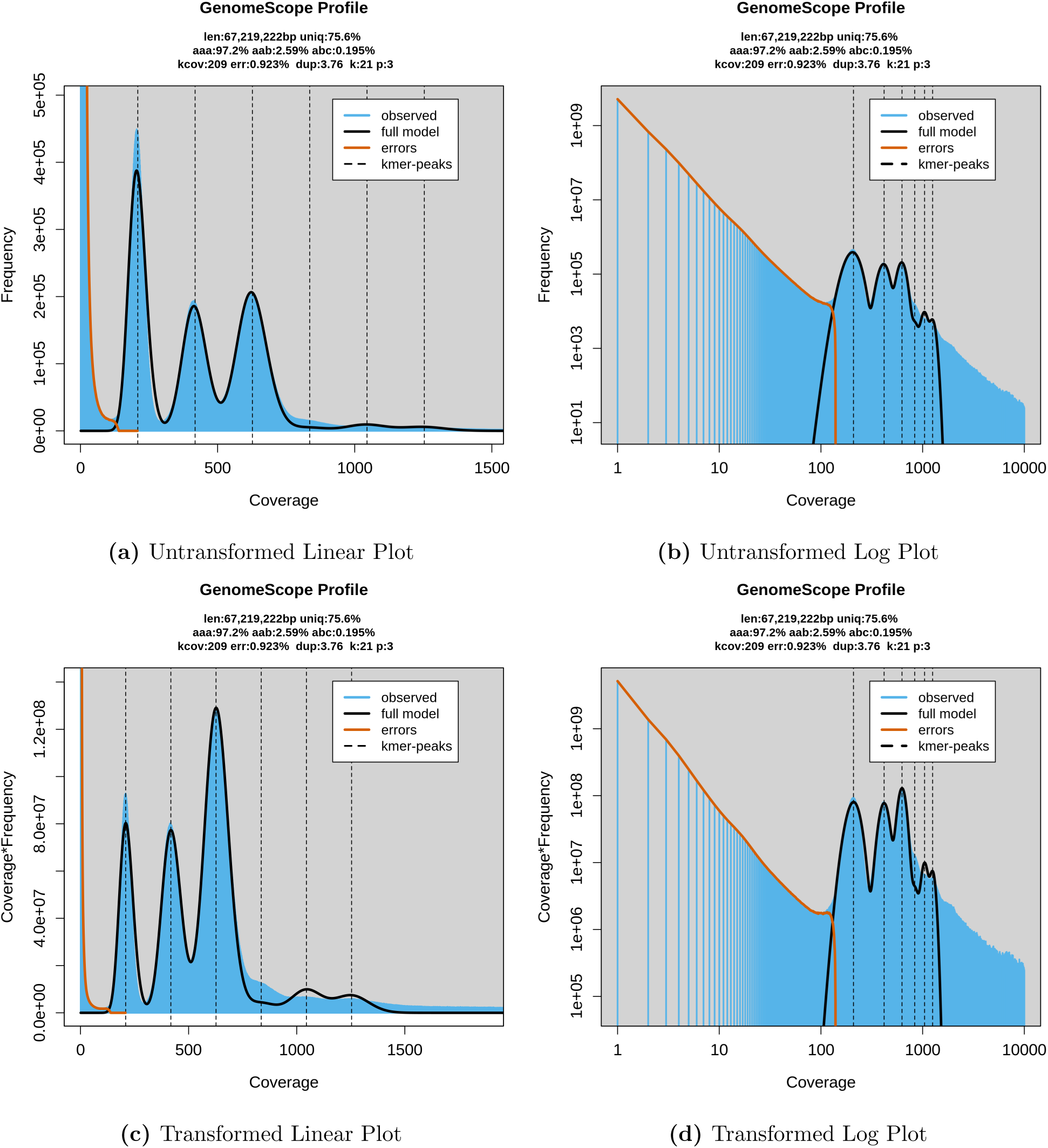
GenomeScope results for *Meloidogyne floridensis*.

**Figure S17:**
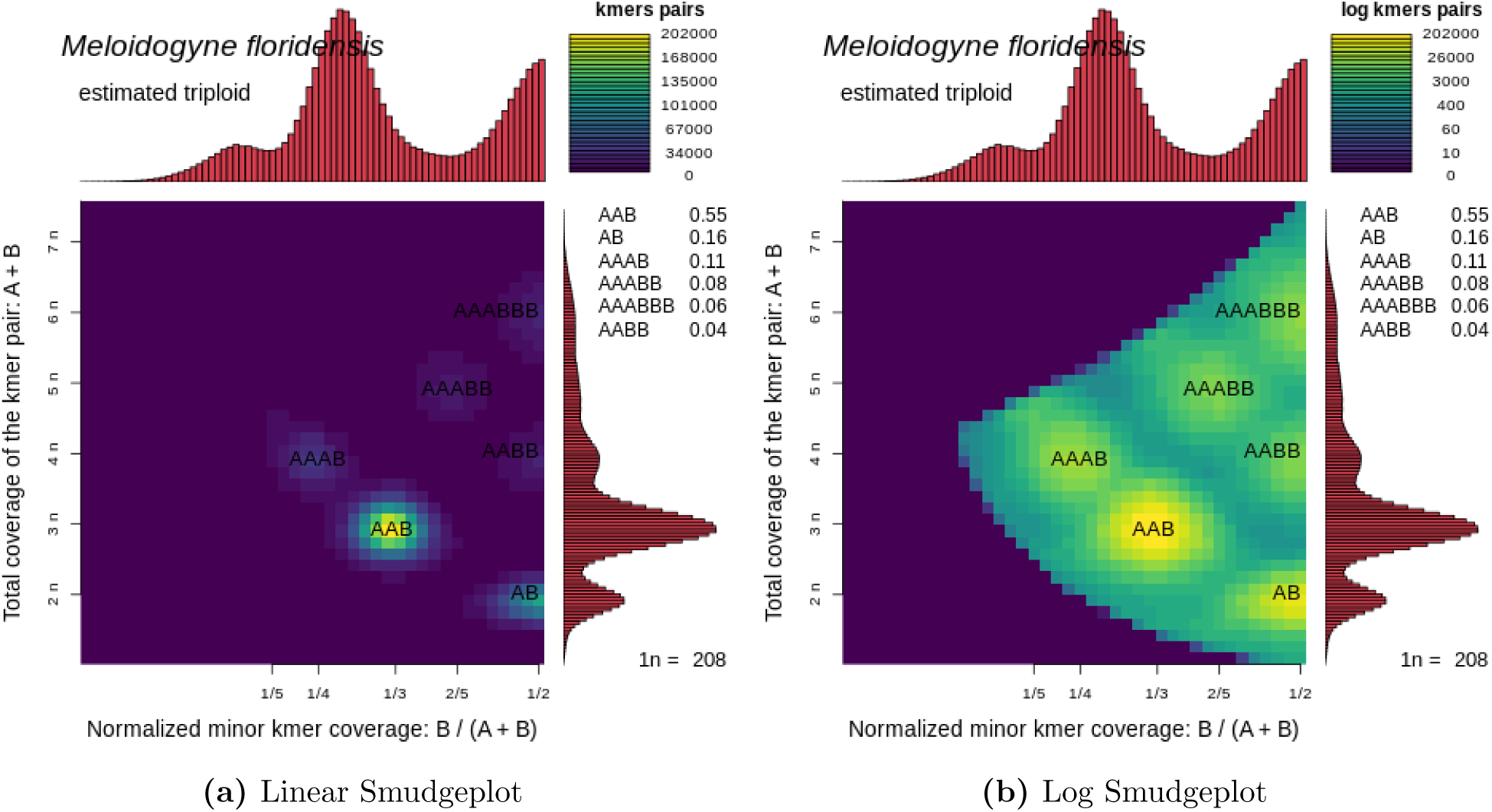
Smudgeplot results for *Meloidogyne floridensis*.

**Figure S18:**
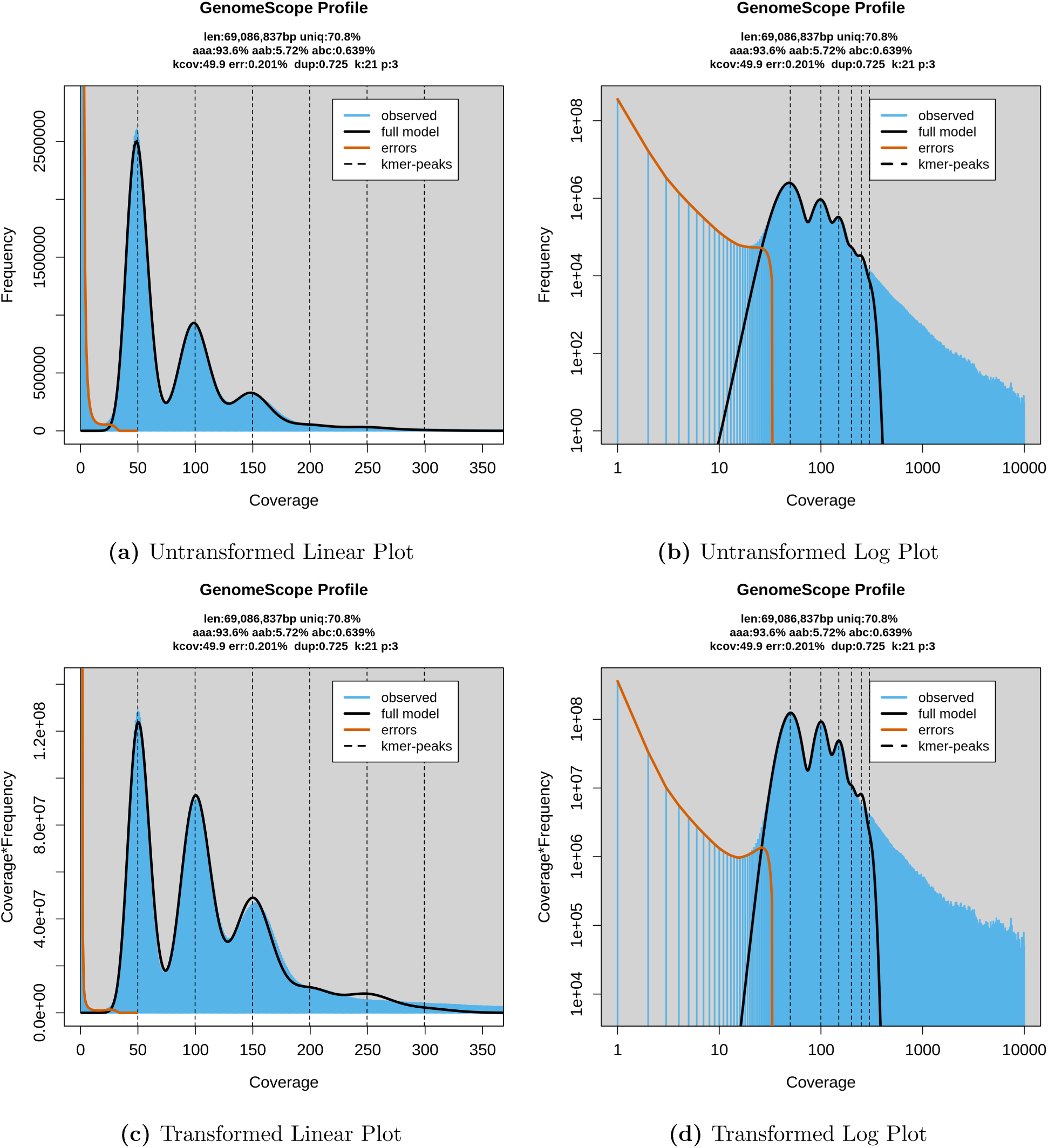
GenomeScope results for *Meloidogyne incognita*.

**Figure S19:**
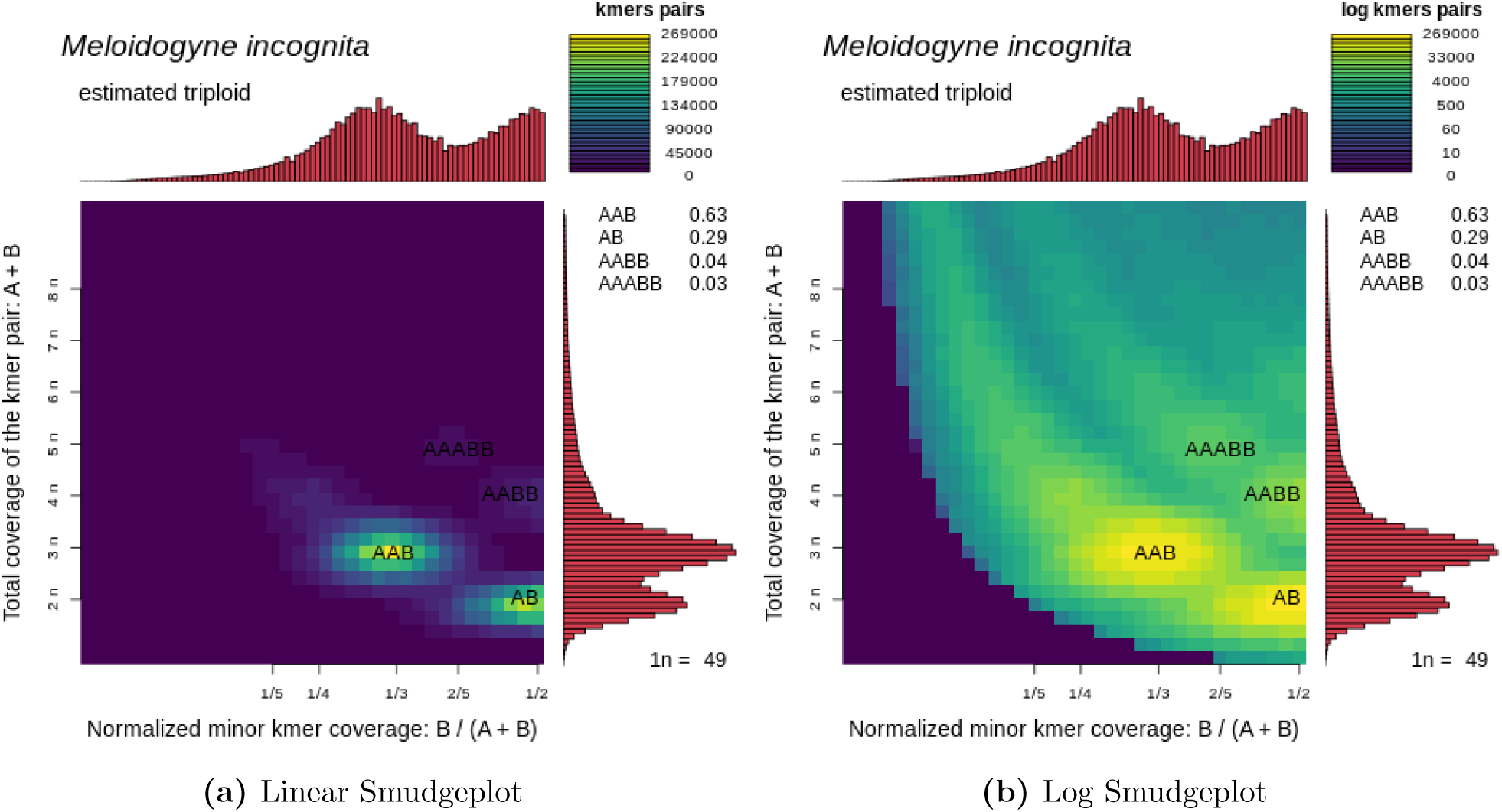
Smudgeplot results for *Meloidogyne incognita*.

**Figure S20:**
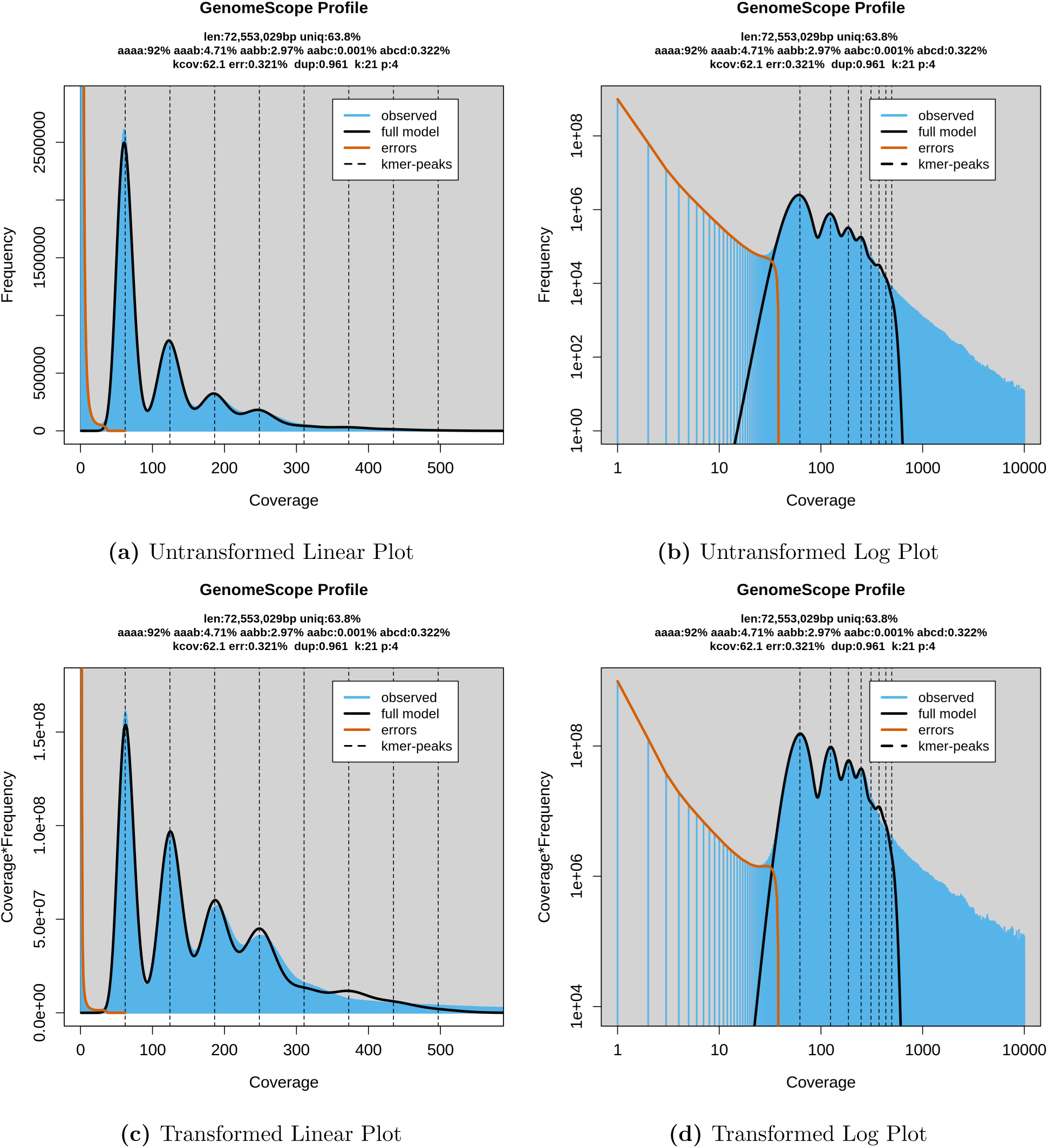
GenomeScope results for *Meloidogyne arenaria*.

**Figure S21:**
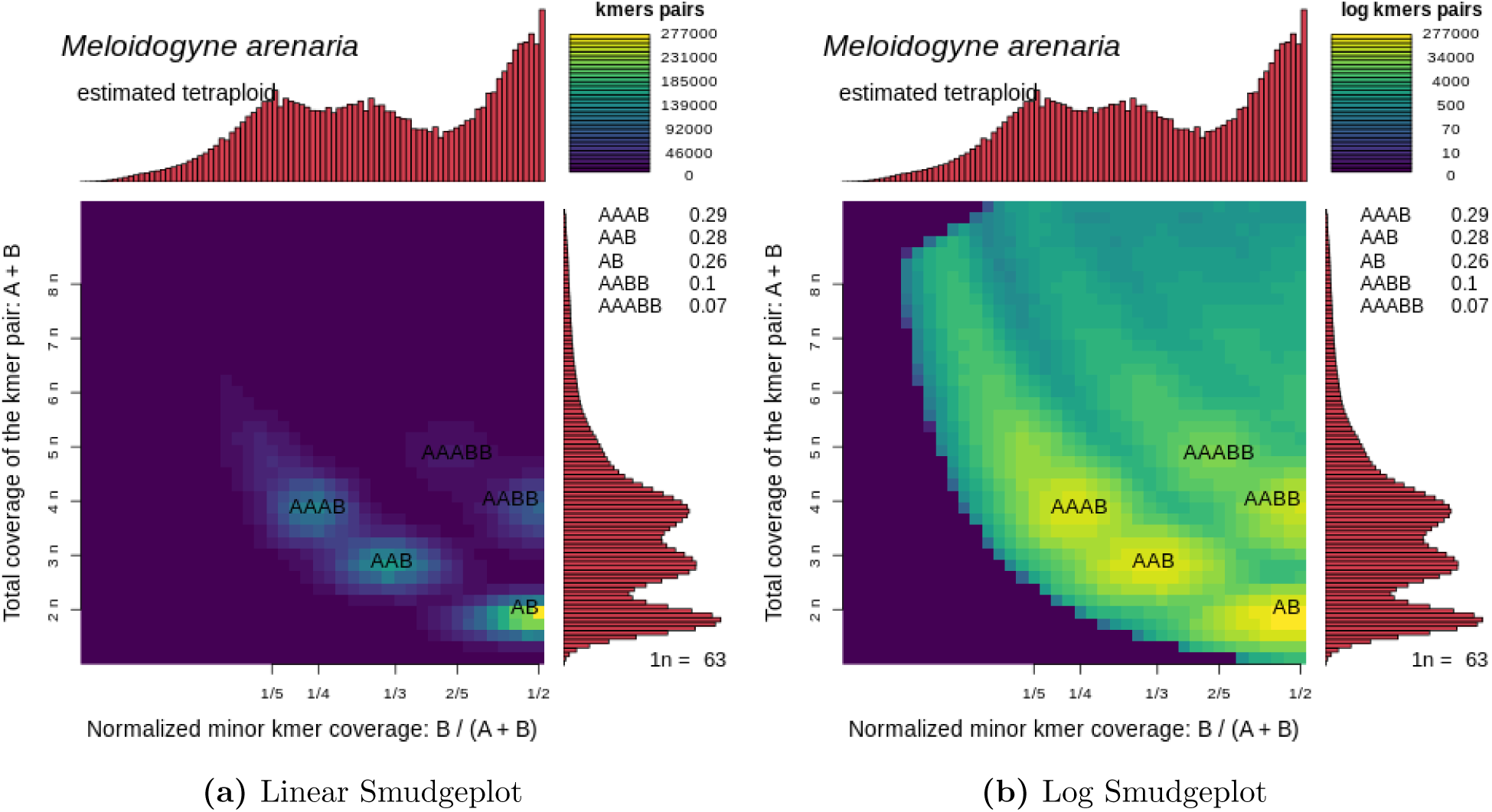
Smudgeplot results for *Meloidogyne arenaria*.

**Figure S22:**
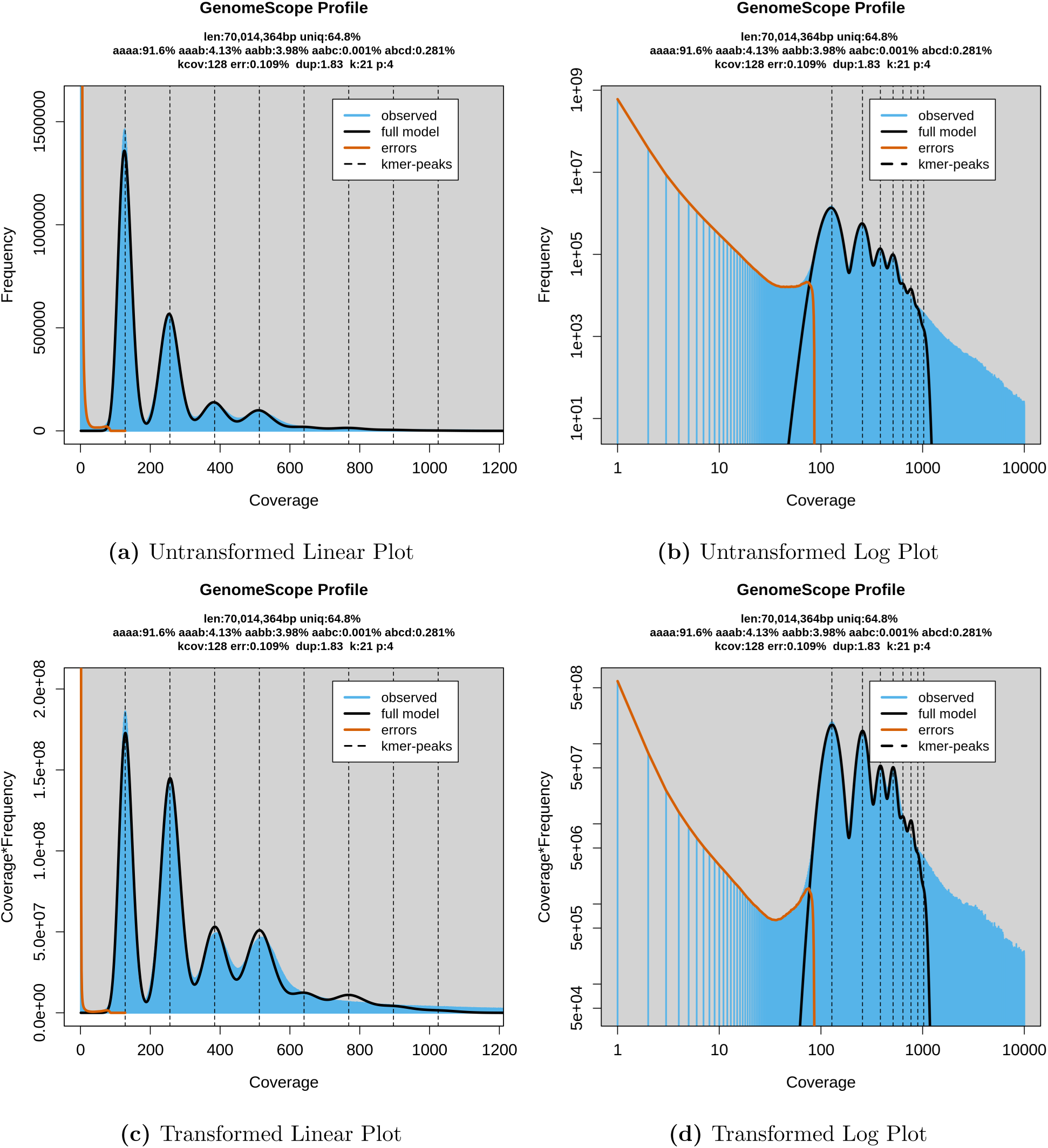
GenomeScope results for *Meloidogyne javanica*.

**Figure S23:**
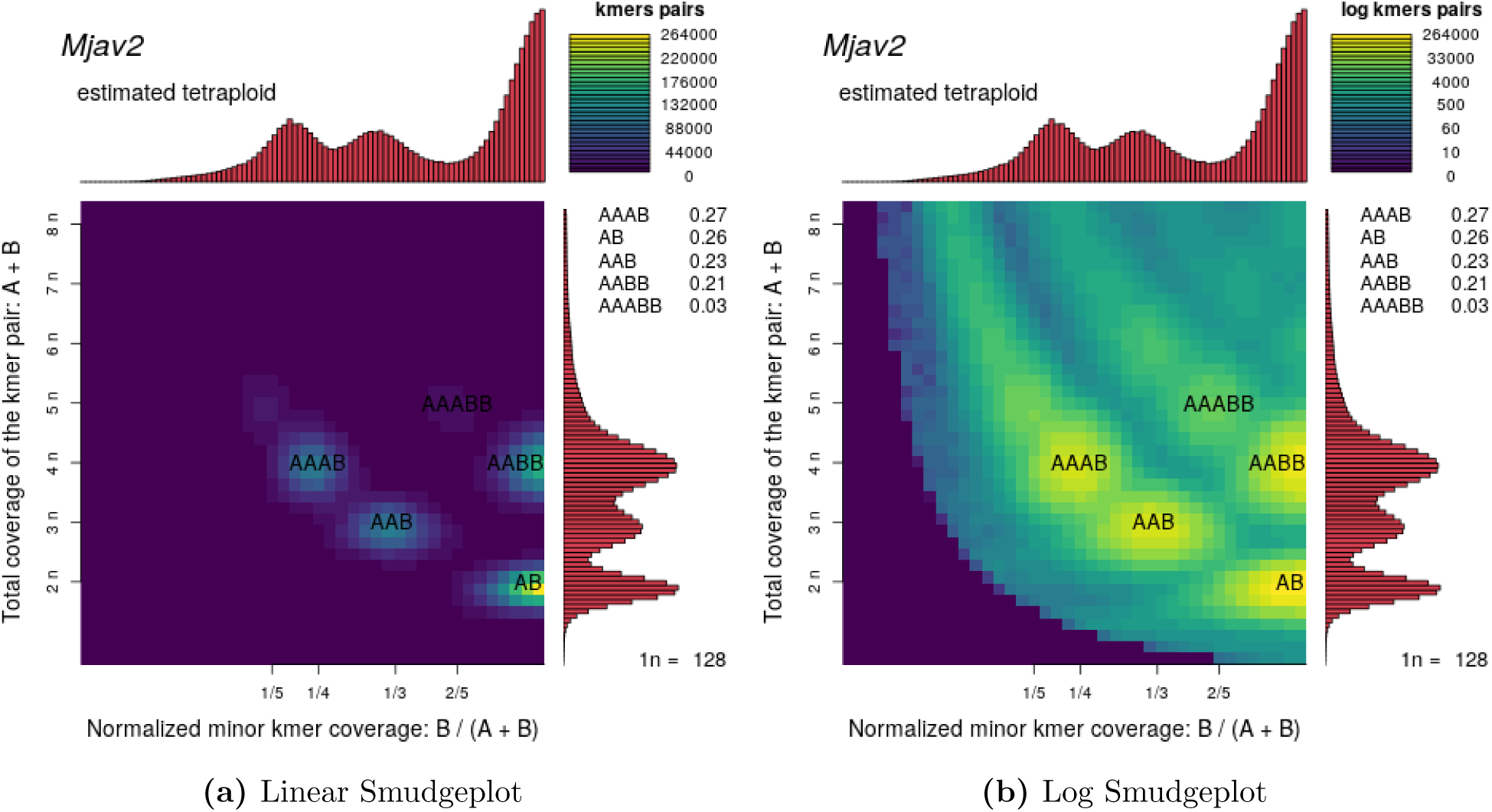
Smudgeplot results for *Meloidogyne javanica*.

### S3.5 Potato Results

**Figure S24:**
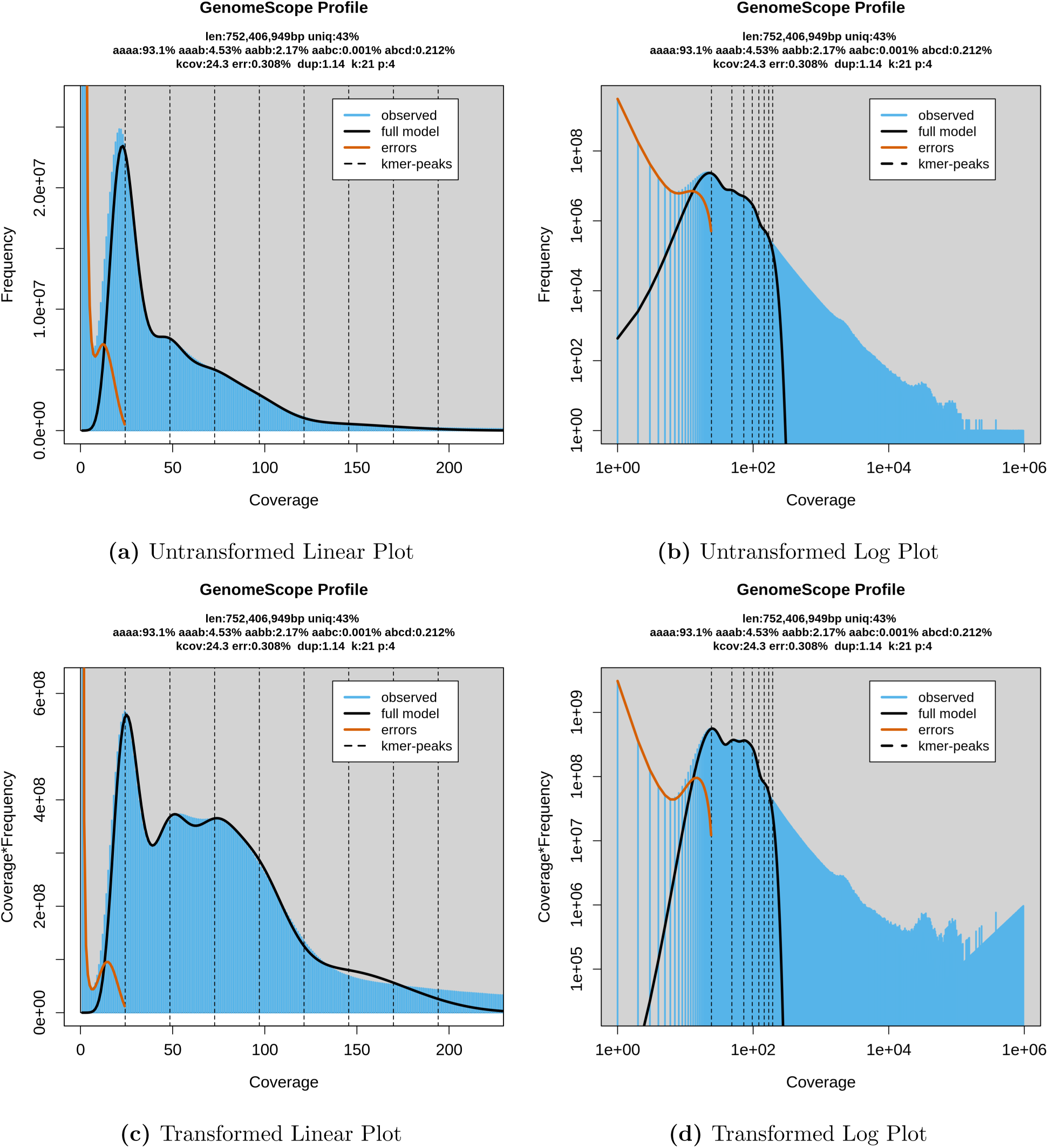
GenomeScope results for *Solanum tuberosum*.

**Figure S25:**
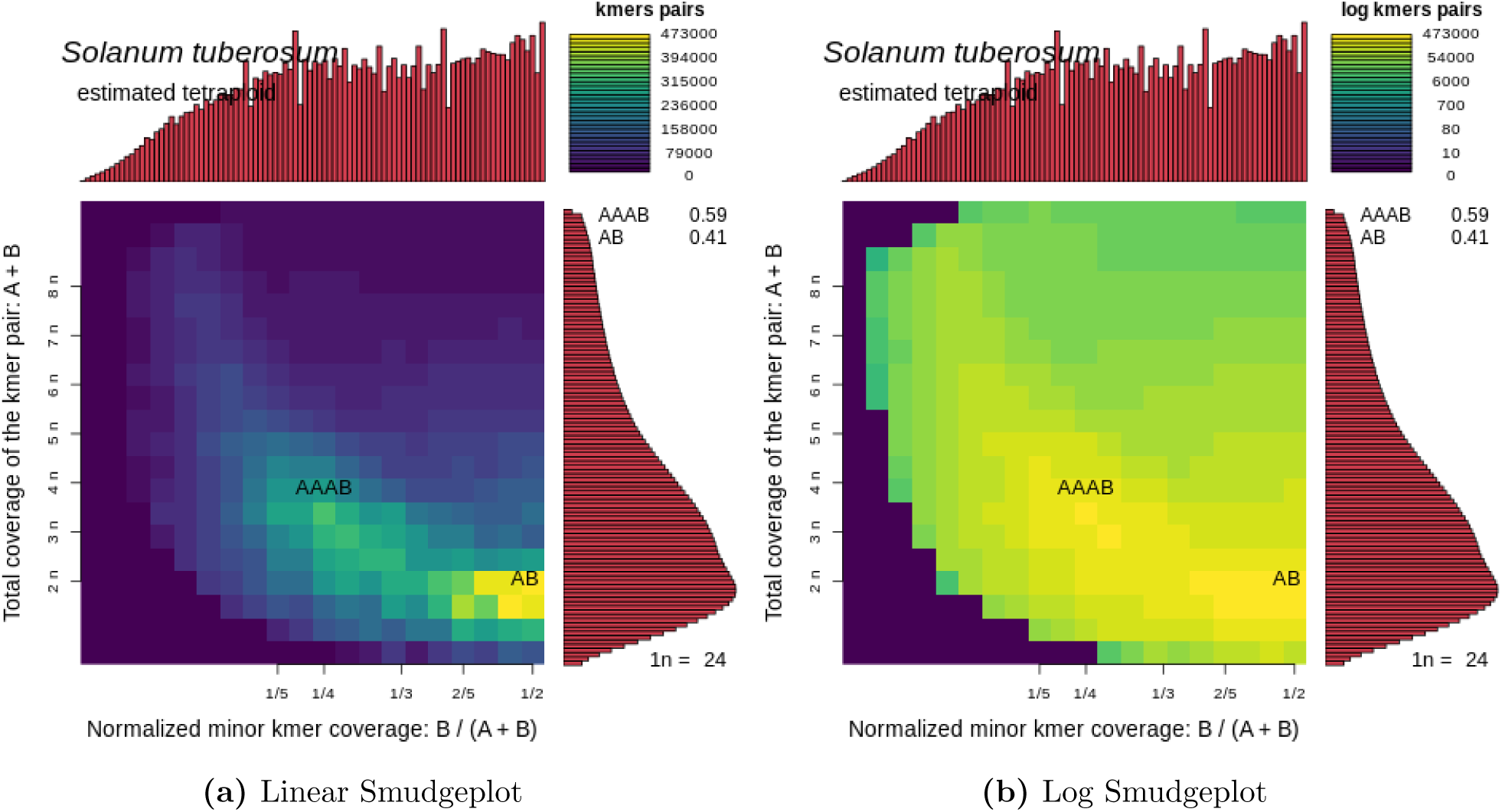
Smudgeplot results for *Solanum tuberosum*.

### S3.6 Wheat Results

**Figure S26:**
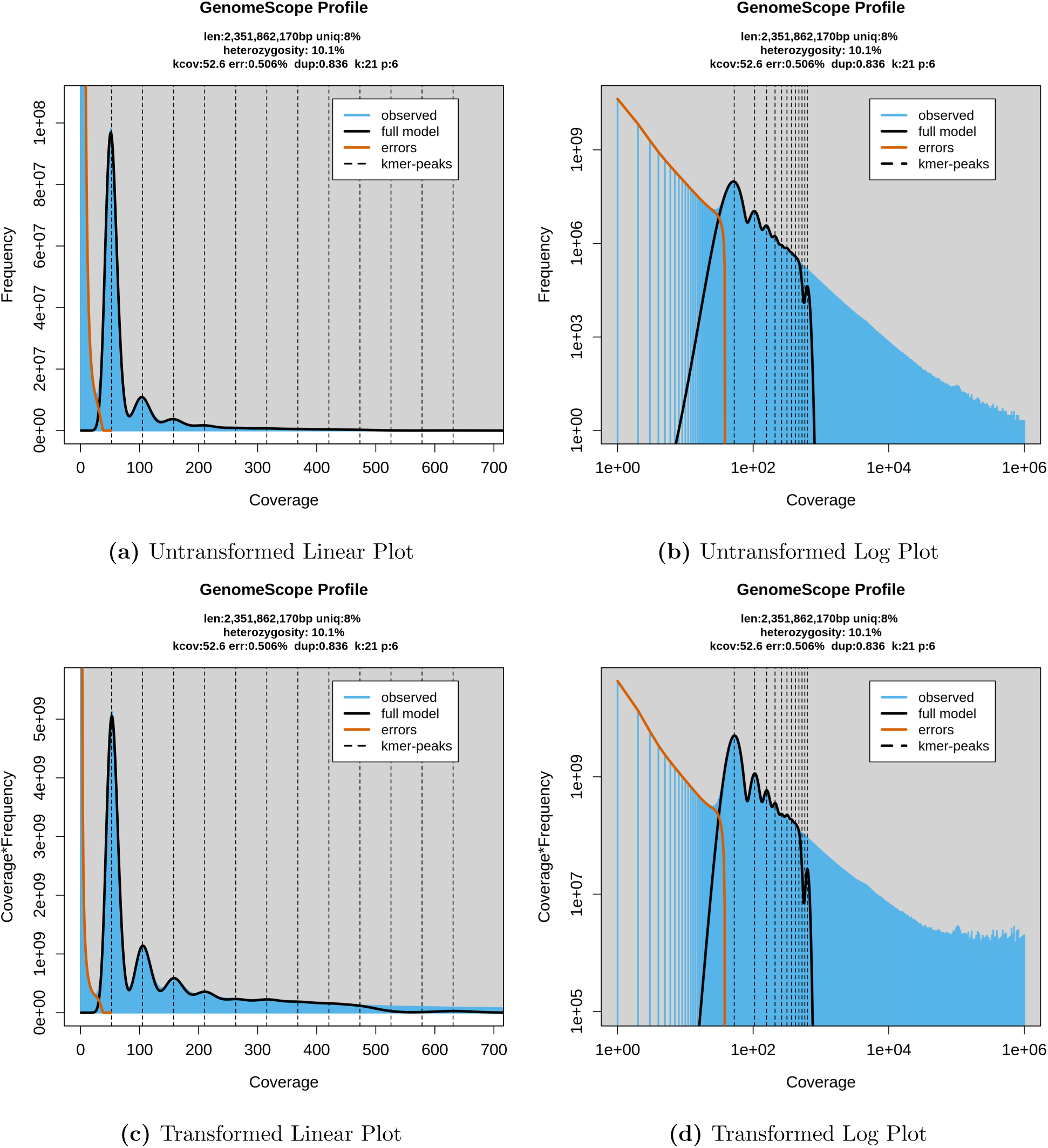
GenomeScope results for *Triticum aestivum*.

**Figure S27:**
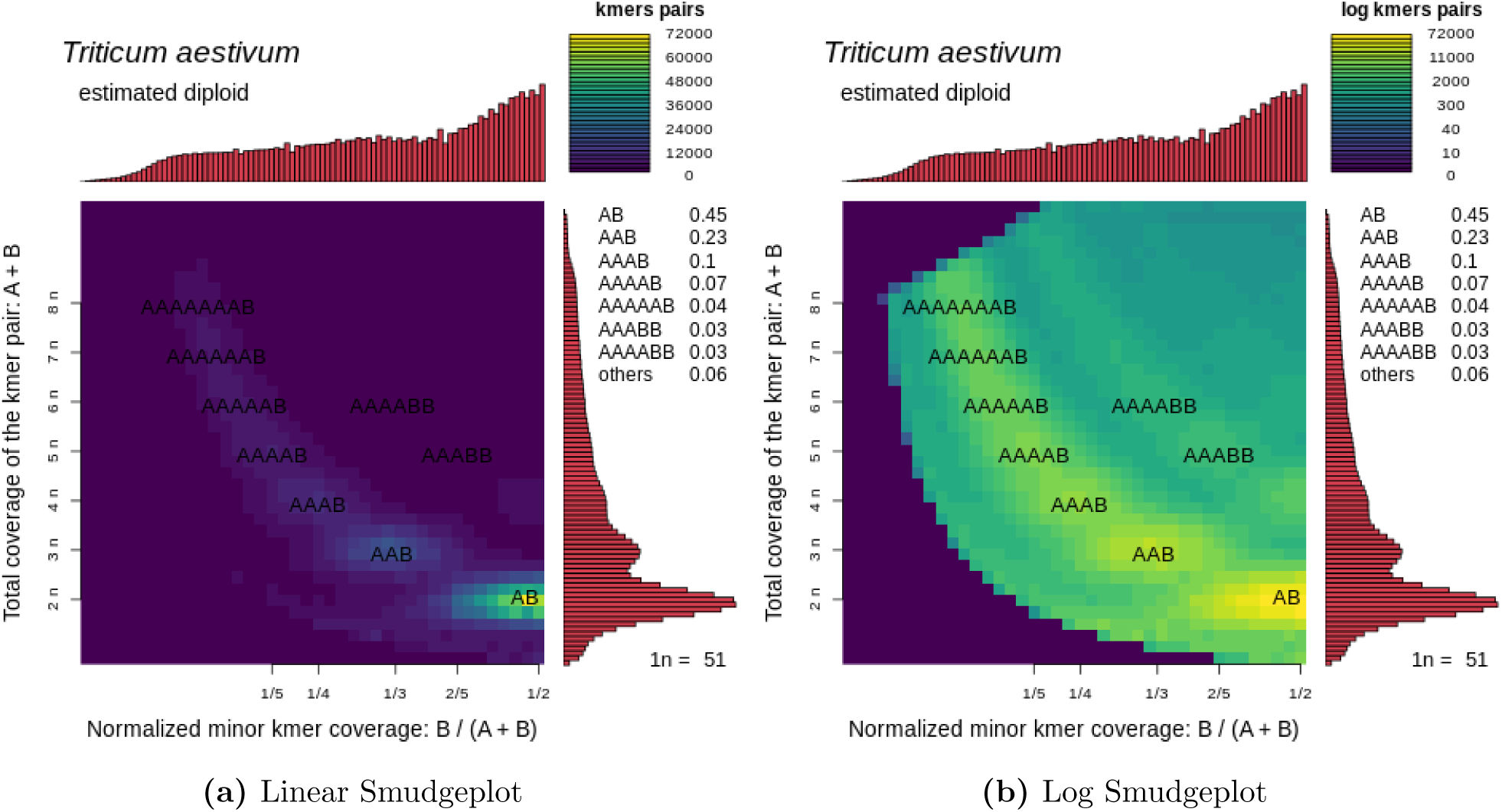
Smudgeplot results for *Triticum aestivum*.

### S3.7 Smudgeplot for Diploid Strawberry Results

**Figure S28:**
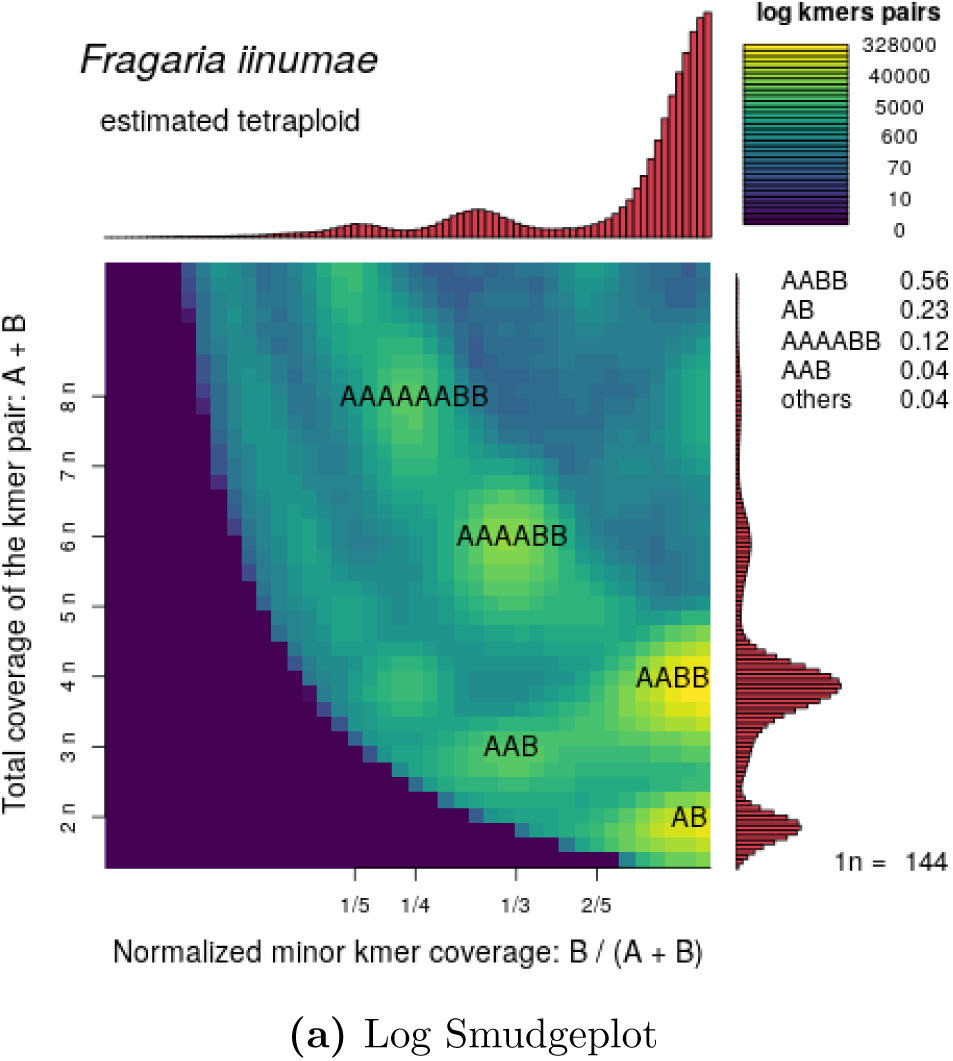
Smudgeplot results for *Fragaria iinumae*.

